# Interference Between Pavlovian Cues and Instrumental Control: A Stratified Comparison Across Alcohol Use Disorder and Smoking

**DOI:** 10.64898/2026.09.25.754324

**Authors:** Louis Thill, Hao Chen, Christian Bäuchl, Samanda Krasniqi, Maren Born, Michael Marxen, Eva Friedel, Maximilian Pilhatsch, Andreas Heinz, Michael N. Smolka

## Abstract

Pavlovian cues can interfere with goal-directed behavior, particularly when they conflict with the required instrumental response. Previous research linked an increased susceptibility to such Pavlovian-to-instrumental transfer (PIT) interference to risky drinking and alcohol dependence. However, it remained unclear whether the observed behavioral and neural alterations generalize to non-treatment-seeking individuals with mild-to-moderate alcohol use disorder (AUD) and to other substances like tobacco. We therefore studied 200 participants from the general population who were stratified by AUD and smoking status before completing a single-lever PIT task during functional magnetic resonance imaging. PIT interference was assessed as the difference in error rates between incongruent PIT trials, in which a Pavlovian conditioned stimulus conflicted with the required instrumental response, and congruent trials. Associations with AUD and smoking status, as well as dimensional associations of PIT interference and related neural activation with substance-related measures, were examined. As expected, participants with AUD showed greater PIT interference and reduced activation in the lateral prefrontal, dorsomedial prefrontal, and anterior cingulate cortices during PIT conflicts than participants without AUD. The magnitude of these behavioral and neural alterations also correlated with higher AUD severity and alcohol consumption, while being independent of smoking status and other smoking-related measures. These findings suggest that enhanced PIT interference generalizes to the broader AUD population but may not extend to smoking. They further implicate a reduced recruitment of prefrontal control networks during PIT conflicts as a potential mechanism underlying enhanced PIT interference in AUD and its association with clinically relevant outcome measures.

## Introduction

Alcohol represents a major public health challenge, contributing substantially to the global burden of disease and causing 2.6 million deaths annually [1]. Worldwide, over 400 million people experience clinically significant impairments or distress due to problematic drinking patterns, thereby meeting the criteria for an alcohol use disorder (AUD) [1, 2]. Despite the considerable health risks and societal costs associated with the development of an AUD, our understanding of the underlying neurocognitive mechanisms remains limited [3–5].

A promising line of research highlights the role of associative learning mechanisms and, in particular, an increased influence of Pavlovian conditioned stimuli (CSs) on goal-directed behavior [6, 7]. Studies investigating the effects of Pavlovian cues on instrumental behavior, also known as Pavlovian-to-instrumental transfer (PIT), consistently observed that appetitive CSs promote instrumental responding and approach, whereas aversive CSs promote inhibition and avoidance [8]. More importantly, several studies found that risky drinkers and alcohol-dependent patients show a stronger invigoration or suppression of instrumental approach than matched controls when appetitive or aversive CSs are concurrently presented [9–11]. In addition to these increased motivational PIT effects, risky drinkers and alcohol-dependent patients were also more likely to show inadequate approach or avoidance when Pavlovian CSs conflicted with the required instrumental response, indicating stronger PIT interference [12, 13]. An increased susceptibility to PIT interference was also linked to increases in alcohol consumption [14] and relapse after detoxification [12].

At the neural level, these elevated PIT effects in risky drinkers and alcohol-dependent patients were associated with dysregulations in a fronto-striatal-limbic network [13, 14]. Relative to controls, risky drinkers and alcohol-dependent patients showed stronger PIT-related neural activation in the ventral striatum (VS) [10, 11, 13] and amygdala [15], which have been implicated in bottom-up motivational processes underlying PIT that may be generally altered in AUD [7, 16–21]. PIT-related neural alterations were also found in the dorsomedial and lateral prefrontal cortex (dmPFC, lPFC), which were associated with PIT interference, risky drinking, and increases in alcohol consumption [13, 14]. Complementary to the VS and amygdala, these regions may reflect an adaptive recruitment of top-down control when Pavlovian CSs interfere with goal-directed behavior [13, 22].

Taken together, these PIT studies indicate that enhanced PIT may be relevant for alcohol consumption, the development of an AUD, and its maintenance, while pointing to alterations in a fronto-striatal-limbic network as neural substrates. However, several research gaps remain. First, previous studies have primarily focused on young risky drinkers and detoxified alcohol-dependent inpatients, representing early and very severe stages of AUD [12, 13]. Consequently, it remains unclear whether the observed behavioral and neural alterations generalize to the large population of non-treatment-seeking individuals with predominantly mild-to-moderate AUD, which often goes clinically undetected [23, 24]. Due to its proximity to the transition from risky drinking to AUD, this population may provide deeper insights into the neurocognitive mechanisms of PIT and how they may vary across the AUD spectrum. Second, despite PIT representing a basic mechanism of action control that may be generally relevant for substance use [25], it is still unknown whether comparable PIT alterations can be found for other substance use disorders (SUDs). The only available studies on human PIT beyond AUD investigated the impact of smoking-related Pavlovian CSs, showing that daily and non-daily smokers exhibit an invigoration of drug-seeking behavior in response to smoking-related cues compared to neutral ones [26–31]. However, these studies did not examine whether smokers are more prone to PIT interference than non-smokers and exhibit similar PIT-related neural alterations as risky drinkers and alcohol-dependent patients. They also rarely studied the relationship between PIT and continuous smoking-related measures, leaving unresolved whether an increased susceptibility to PIT correlates with greater tobacco consumption and tobacco use disorder (TUD) severity. A study by Hogarth and Chase [30] even indicates that the magnitude of smoking-specific PIT may be independent of smoking severity, thereby challenging the account of enhanced PIT representing a shared mechanism across SUDs [30]. Despite these open questions, PIT studies in risky drinkers and alcohol-dependent patients did not control for smoking status, making it unclear to what extent the reported PIT effects were attributable to AUD, smoking, or their frequent co-occurrence [32].

To address these gaps, we conducted a longitudinal study of PIT interference in a general population sample comprising non-treatment-seeking individuals with predominantly mild-to-moderate AUD and individuals without AUD. Smoking status was balanced within each group by stratifying participants into non-smokers, occasional smokers, and daily smokers during recruitment. Based on the existing literature, we expected stronger PIT interference in individuals with AUD than in those without. Consistent with the hypothesized transdiagnostic relevance of PIT [6, 7], we also predicted a systematic increase in PIT interference with increasing tobacco use, resulting in maximally elevated PIT in daily smokers with AUD. At the neural level, we expected participants with AUD, relative to those without, to show stronger PIT-related brain responses in the VS and amygdala, reflecting stronger bottom-up motivational processes, and weaker brain responses in the lPFC and dmPFC, indicating reduced top-down control. The same pattern of neural alterations was also predicted for smokers, with their magnitude increasing with increasing smoking status. In addition to these group effects, we anticipated dimensional associations of PIT interference and the hypothesized neural alterations with continuous substance-related measures, particularly with higher AUD and TUD severity, higher consumption quantity, and higher consumption frequency.

## Methods

The study was part of the TRR265 “Losing and Regaining Control over Drug Intake” and was preregistered at ClinicalTrials.gov (ID NCT05992272). All procedures complied with the Declaration of Helsinki, were approved by the ethics committee of Technische Universität Dresden (EK 512122018), and were explained to the participants, who gave full written consent.

### Recruitment and Study Design

Participants were recruited in Dresden, Germany, using flyers, online advertisements, and other public postings. Eligibility criteria included age between 18 and 65 years, normal or corrected-to-normal vision, right-handedness, and proficiency in written and spoken German. Any previous history of a neurological or major psychiatric disorder (except AUD, TUD, and cannabis use disorders), inpatient treatment for any SUD, severe head trauma, contraindication for functional magnetic resonance imaging (fMRI), or a recent intake of central nervous system agents (within the last 10 days or fewer than five half-lives before assessment) led to immediate exclusion. Candidates with severe withdrawal symptoms or the intention to seek treatment for any SUD were also excluded.

Candidates meeting these criteria were invited to a baseline assessment, which comprised sociodemographic data collection, structured clinical interviews for DSM-5 (AUD, TUD, other selected disorders) [33], neuropsychological tests, and questionnaires. The latter included two quantity-frequency questionnaires for alcohol use (QFQ-A) [34] and smoking (QFQ-S; items provided in Supplemenatry Material S1), the AUD Identification Test (AUDIT) [35], the Fagerström Test for Nicotine Dependence (FTND) [36], the Craving Automated Scale for Alcohol (CAS-A) [37], and the Questionnaire on Smoking Urges (QSU-G) [38].

Based on these measures, AUD and smoking status were balanced at baseline by stratifying participants into a 2 x 3 design. Participants meeting more than one DSM-5 criterion for AUD were classified as ‘AUD’, whereas participants meeting fewer criteria were classified as ‘non-AUD’. Each participant was also categorized by the number of smoking days in the last three months before the study, with participants reporting daily smoking being classified as ‘daily smokers’, participants reporting more than one but fewer than 90 smoking days as ‘occasional smokers’, and participants reporting no smoking as ‘non-smokers’.

All participants included in this study also completed an MRI session (within 4 weeks), during which they performed an established single-lever PIT task [39]. They were also invited to three follow-ups over 12 months on substance use-related measures.

For each session, participants were instructed to abstain from drinking (*≥* 24*h*), smoking (*≥* 1*h*), and other drug use (*≥* 24*h*), and abstinence was verified via breathalyzer and urine-based drug screening.

### PIT Paradigm

PIT interference was assessed using a four-part single-lever PIT paradigm [39], which demonstrated excellent split-half reliability (*r* = .95–.97) and an acceptable 3-week test-retest reliability (ICC = .53) in a clinical sample [40]. The employed joystick-based version was programmed in MATLAB using the Psychophysics Toolbox Version 3 [41–43] and provided a balanced approach-avoidance design optimized for functional imaging [39].

Figure 1 illustrates the first three parts of the paradigm, which constitute the core elements for assessing PIT interference. These comprised an instrumental learning (Part 1), a Pavlovian conditioning (Part 2), and a PIT phase (Part 3). During Parts 1 and 2, participants learned the action-outcome associations for six different shells (i.e., instrumental stimuli) and the stimulus-outcome associations for 5 different audiovisual compound cues (i.e., Pavlovian CSs). In Part 3, they repeated the instrumental approach-avoidance task from Part 1 in the MRI scanner while Pavlovian CSs from Part 2 were presented concurrently. Participants were instructed to maximize their monetary gain during Part 3 by collecting ‘good’ shells, whose collection was associated with monetary gains during Part 1, and rejecting ‘bad’ shells, whose collection was punished by monetary losses, without receiving feedback about the correctness of their actions.

**Figure 1.**
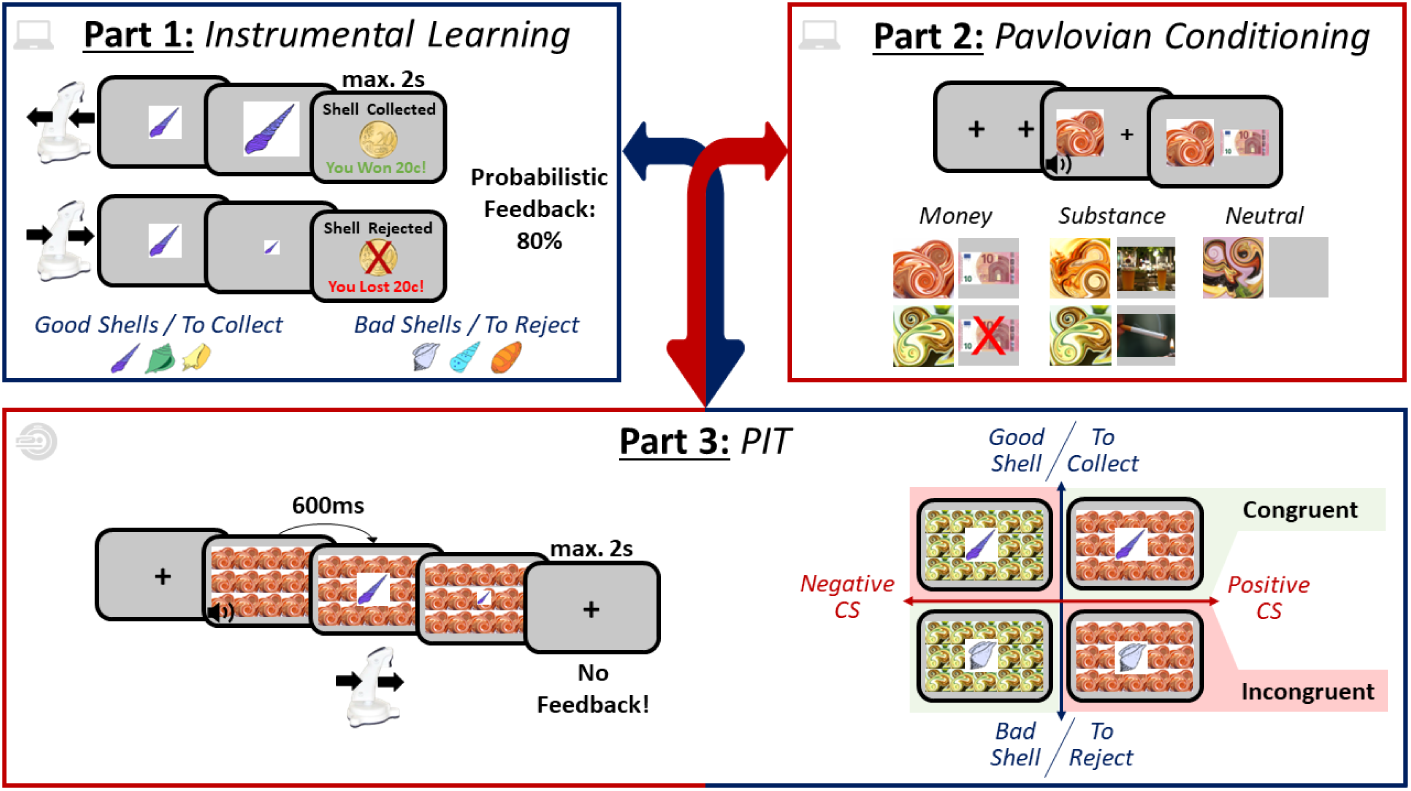
Experimental Design of the Single-Lever PIT Paradigm. *Note.* The figure is based on the task schematic from Belanger et al. [39], with each box corresponding to one of the first three parts of the PIT paradigm. In *Part 1*, participants learned the action-outcome associations for six different shells through trial and error. Each shell was unique in its shape and color and was randomly assigned to be either ‘good’ or ‘bad’ (counterbalanced). Participants were instructed to collect ‘good’ shells by pulling (i.e., approach) and reject ‘bad’ shells by pushing a joystick (i.e., avoidance), while receiving probabilistic feedback. Correct responses were rewarded with 20 cents in 80% of the trials and punished with a loss of 20 cents in 20% of the trials. For incorrect actions, the probabilistic feedback was inverted. Participants were required to complete at least 60 trials in Part 1 and reach a minimum accuracy of 80% across 16 consecutive trials or complete a total of 120 trials. In *Part 2*, participants learned the stimulus-outcome associations for 5 different audiovisual compound cues. Each compound cue consisted of an abstract fractal-like image paired with a musical instrument tone, and was randomly assigned to be associated with monetary gain (+10€), monetary loss (−10€), alcohol, smoking, or a neutral outcome. Participants were instructed to memorize the stimulus-outcome associations by passively observing 16 sequential presentations of the audiovisual compound cues and a visual illustration of their outcomes. This resulted in a total of 80 trials in Part 2. In *Part 3*, participants completed the instrumental task from Part 1 while simultaneously being confronted with the Pavlovian CSs from Part 2. In each trial, one of the five fractal-like images was tiled in the background of one of the shells, and the associated sound was presented through headphones at Pavlovian CS onset. The Pavlovian CSs preceded the onset of the instrumental shell by 600 ms. Participants were instructed to maximize their monetary gain by correctly collecting good shells while rejecting bad ones. Participants completed a total of 180 trials, equally distributed across all 2×5 conditions (shell valence x Pavlovian CS), and conducted under nominal extinction. The trial sequence for each part was randomized for each participant.

In addition to these parts, participants completed a set of query trials (Part 4) and a debriefing questionnaire. During the query trials, participants were forced to choose between pairs of Pavlovian CSs to assess whether they had learned the relative valence of the CSs (for more details, see Supplementary Material S2). Complementarily, the debriefing questionnaire assessed participants’ general understanding of the task by asking them to classify a set of task-related statements and rate cue- and outcome-related features.

For a more detailed description of the task, see Belanger et al. [39].

### Continuous Substance-Related Measures

AUD and TUD severity were assessed via AUDIT and FTND total scores, whereas substance-specific consumption was derived from QFQ-A and QFQ-S. Drinking and smoking quantity was computed as the average daily intake of alcohol and tobacco (g/day and units/day, respectively), whereas drinking and smoking frequency were defined as the number of drinking and smoking days per week.

For exploratory analyses, we also computed the CAS-A total score, the QSU-G mean score, both assessed at baseline, and the AUDIT-Consumption subscore (AUDIT-C) [35] for each time point. AUDIT-C was included as an integrative measure of drinking across both quantity and frequency to investigate the association between greater PIT interference and drinking trajectories [cf. 14]. CAS-A and QSU-G were used to assess substance-specific craving, which has been identified as a clinically relevant outcome measure associated with cue effects, substance use, and relapse, particularly in AUD [44–47].

### MRI Acquisition and Preprocessing

All MRI data were acquired on a 3T Siemens MAGNETOM® Tim Trio (Siemens Healthineers AG, Forchheim, Germany) with a 32-channel head coil and preprocessed with SPM12 (Wellcome Department of Cognitive Neurology, Institute of Neurology, London, United Kingdom). Detailed information about acquisition parameters and preprocessing is provided in the Supplementary Material S3.

### Statistical Analysis

All statistical analyses, if not stated otherwise, were conducted with R 4.4.4. (R Core Team, Vienna, Austria) and considered significant at a threshold of *p* < .05 (uncorrected). One-tailed tests were used for AUD- and smoking status-related analyses targeting the predefined hypotheses.

#### Behavioral Data

##### Computation of PIT Interference

First, the raw joystick data was preprocessed and an initial task manipulation check was conducted (see Supplementary Material S4). Then, PIT trials were classified by trial type (substance-vs. money-related Pavlovian CS) and congruity. Money-related trials were classified as ‘congruent’ when a ‘good’ shell was preceded by an appetitive Pavlovian CS (+10€) or a ‘bad’ shell was preceded by an aversive CS (−10€). Trials with opposite valences of Pavlovian CS and instrumental shell were classified as ‘incongruent’. For each participant, Δ*ER* was then computed as the primary outcome for PIT interference by subtracting the error rate (ER) across congruent PIT trials from the ER across incongruent PIT trials [cf. 13, 39].

##### Association With AUD and Smoking Status

To investigate the hypothesized group differences, robust linear regressions (rlm) were fitted using the *MASS* package [48]. This approach was selected for its robustness to violations of distributional model assumptions, including the observed non-normality of Δ*ER* and its residuals, while retaining the ability to include covariate and interaction effects [49]. Δ*ER* was modeled as a function of AUD (−0.5 = ‘Non-AUD’, 0.5 = ‘AUD’), smoking status (mean-centered), and their interaction. We also fitted an extended version of this model, including age and gender as additional regressors, to assess potential covariate effects. For both models, *p*-values and 95% bias-corrected and accelerated confidence intervals (CIs) were obtained via bootstrapping with 5000 resamples, using the *boot.pval* package.

##### Associations With Continuous Substance-Related Measures

As the previous analysis provided no evidence for significant age or gender effects on PIT interference, the hypothesized bivariate associations with AUD and TUD severity, alcohol and tobacco consumption, and substance-specific craving were assessed using one-sided Kendall’s tau rank correlation coefficients (*τ*). Similar to rlm, this non-parametric approach provided robust estimates when distributional assumptions were violated, including non-normality, outliers, and tied ranks, as observed in the data [50]. To assess whether PIT interference predicted drinking trajectories across time points, a robust linear mixed-effects model (rlmer) was fitted using the robustlmm package. This model included AUDIT-C scores as dependent variables and PIT interference, the separate time points, and their interaction as fixed effects. Random intercepts at the participant level were included to account for inter-individual differences. 95% CIs and *p*-values were assessed for all fixed effects via participant-level bootstrapping with 1000 resamples.

To facilitate the comparison with the broader PIT literature and provide a more comprehensive characterization of non-drug-related PIT, we also conducted an exploratory analysis of motivational PIT in the present sample (provided in Supplementary Material S5).

### MRI Data

The individual brain responses during the PIT phase were analyzed using general linear models (GLMs) specified for each participant and convolved with the canonical hemodynamic response function [51]. The GLMs comprised 10 regressors of interest corresponding to the 2×5 conditions (shell valence x Pavlovian CS) as well as 25 nuisance regressors. The latter were included to account for higher-order head motion and comprised six rigid-body motion parameters estimated during spatial realignment, 18 additional motion regressors derived from a Volterra expansion [52], and a motion censoring regressor excluding volumes with a framewise displacement > 0.9 [53]. To control for low-frequency noise and temporal autocorrelation, the functional images were high-pass filtered with a cutoff of 128s and pre-whitened using SPM12’s FAST model.

In the next step, two first-level contrasts were created for each subject. The first contrast was designed to mirror the behavioral PIT interference measure by contrasting the incongruent and congruent conditions (i.e., neural PIT interference). The second contrast was based on a recent reliability investigation [40], which identified the contrast between the incongruent condition and baseline as the more reliable and promising contrast for predicting individual outcomes (i.e., neural incongruity). Notably, this contrast has already been used by previous studies that linked neural activation during incongruent PIT trials to risky drinking and subsequent drinking trajectories [13, 14].

To test the predefined hypotheses regarding lPFC, dmPFC, VS, and amygdala activation during PIT, we then defined four regions of interest (ROIs; see Figure 2) and extracted the mean beta-weights for each from the neural PIT interference and neural incongruity contrasts. In a first step, we tested for increased activation in the lPFC and dmPFC during incongruent vs. congruent PIT trials using one-sided one-sample *t*-tests, and for a direct association between behavioral and neural PIT interference in each ROI using Kendall’s rank-based correlations. In the next step, we inspected the hypothesized AUD group differences in neural PIT interference and neural incongruity for each ROI by performing one-sided *t*-tests on the extracted beta weights. For completeness, we also tested for smoking status-related and AUD x smoking status interaction effects on neural PIT interference and neural incongruity using linear regression models (lm). Lastly, Pearson correlations were computed to examine the associations between increased VS and amygdala activation during incongruent PIT trials, reduced lPFC and dmPFC activation, and greater substance-related measures. To account for multiple testing within test families, *p*-values from the ROI analysis were False Discovery Rate (FDR)-corrected.

**Figure 2.**
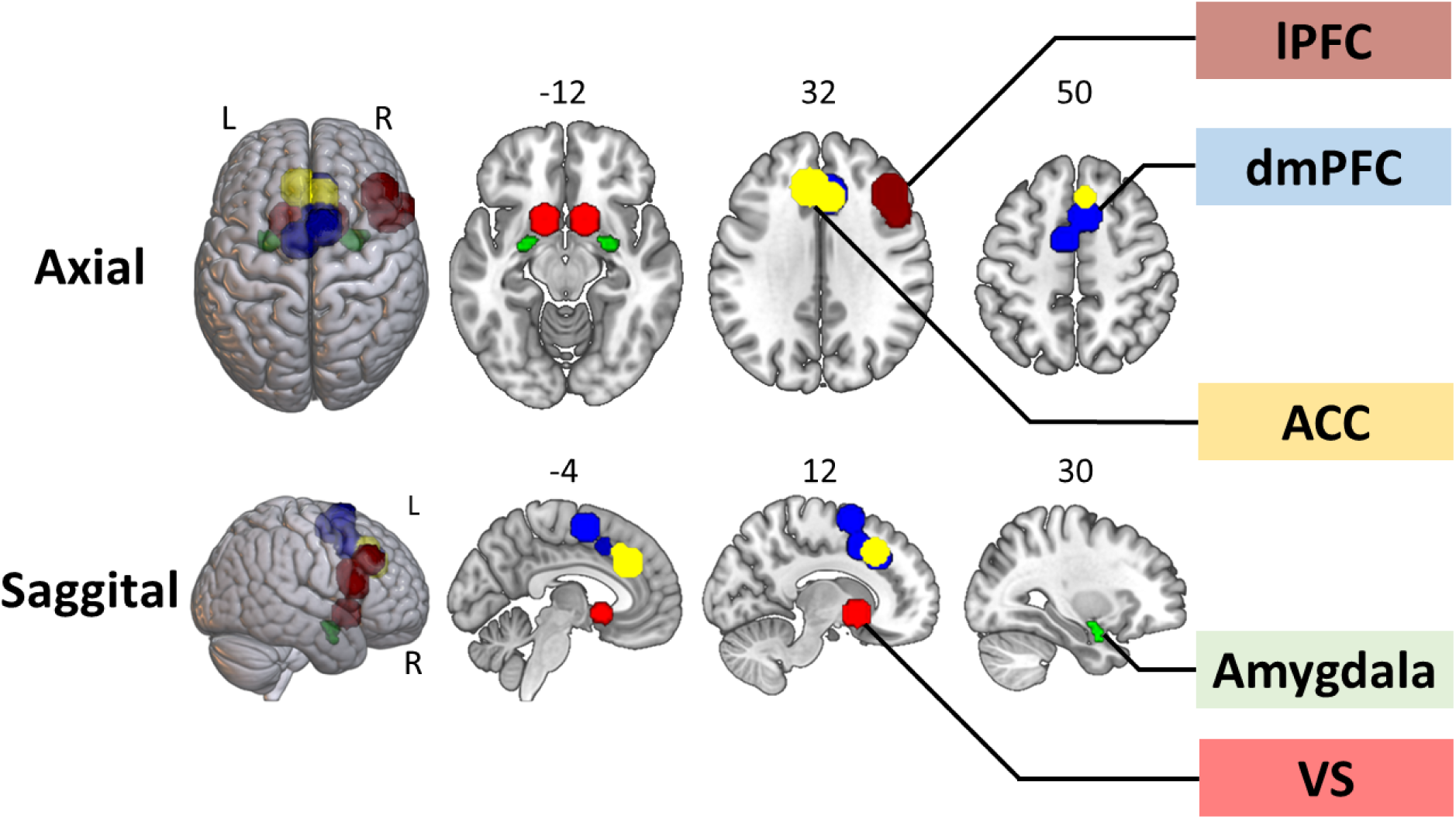
ROI Masks. *Note.* The ROIs comprised four predefined ROIs within the right lPFC, bilateral dmPFC, bilateral VS, and bilateral amygdala [as in 13], as well as an exploratory ROI within the bilateral ACC. Except for the amygdala, all ROI masks were generated as conjunctions of 12-mm spheres centered on meta-analytically derived activation peaks. The right lPFC mask was based on three activation peaks identified by a meta-analysis of Hung et al. [22] on cognitive inhibition (Talairach coordinates: 42/26/30; 46/14/22; 52/16/14). The same meta-analysis was also used to define the dmPFC mask, which included four subregions (Talairach coordinates: 6/14/40; 6/26/32; 8/8/58; −6/0/54). To increase anatomical specificity, the exploratory ACC ROI was defined independently of the dmPFC ROI using two activation peaks in the left and right ACC. These were derived from a meta-analysis by Zhang and Peng [77] on cognitive functions underlying behavioral regulation, including conflict monitoring and inhibitory control (MNI coordinates: −6/30/32; 6/24/40). The bilateral VS mask was based on a meta-analysis by Liu et al. [78], identifying two reward-related activation peaks within the VS (MNI Coordinates: −12/10/-6; 12/10/-6). In contrast to the other ROIs, the amygdala mask was defined anatomically using the AAL atlas in the WFU PickAtlas toolbox [79]. The WFU PickAtlas was also used to convert all activation peak coordinates originally reported in Talairach space to the MNI space.

In addition to these primary analyses, two sets of exploratory analyses were conducted. First, the role of the ACC during PIT conflicts was examined by repeating the same ROI analysis as for the dmPFC. The ACC has consistently been linked to conflict monitoring and the adaptive recruitment of top-down control [54–56] and may therefore be particularly important during PIT when Pavlovian CSs conflict with the required instrumental response. In the present and previous studies [13, 14], the ACC was partially covered by the dmPFC mask, leaving unresolved whether the observed conflict-related effects in this region can be localized more precisely to the ACC. Second, an exploratory whole-brain analysis of neural PIT interference was conducted to identify additional brain regions that showed greater or reduced activation during incongruent, relative to congruent, PIT trials. To this end, individual contrast images were entered into a second-level random-effects analysis and analyzed using one-sample *t*-tests, with *p*(uncorrected) < .001 and a minimum cluster extent of k > 50 voxels.

### Total and Partial Exclusions

Out of the 273 participants who completed both the baseline assessment and the PIT task, 73 were excluded during quality control procedures. 32 were excluded because of technical issues during task administration, 2 because they fell asleep during the PIT phase, and 39 because their behavioral data indicated insufficient task understanding (more details in Supplementary Material S2).

In addition to these hard exclusions, 20 participants were excluded from the MRI analysis only: 17 participants completed the PIT task outside the scanner, and the MRI data for 3 could not be preprocessed.

## Results

### Sample Characteristics

The final sample comprised 200 participants (41.0% female) with a mean age of 31.5 years (*SD* = 10.0, range: 18-63) and generally high educational levels (74% had completed vocational training, A-levels, or higher qualifications).

As summarized in Table 1, 100 participants were classified as ‘AUD’ and 100 participants as ‘non-AUD’. Orthogonally, 70 participants were classified as ‘non-smokers’, 72 as ‘occasional smokers’, and 58 as ‘daily smokers’. Within each stratification factor, groups did not differ in size, gender distribution, or educational level. Only for age, a small but statistically significant difference was observed between the AUD group (*M* = 33.2, *SD* = 11.1) and non-AUD group (*M* = 29.8, *SD* = 8.4; *p* = .015), but not between smoking groups.

**Table 1.** Descriptive Statistics.

|  | **Factor 1: AUD** |  |  |  | **Factor 2: Smoking Status** |  |  |  |  |
|---|---|---|---|---|---|---|---|---|---|
|  | Non-AUD | AUD | Group Difference |  | Non-Smoker | Occasional Smoker | Daily Smoker | Group Difference |  |
|  |  |  | Test | *p* |  |  |  | Test | *p* |
| **Sociodemographic Characteristics** |  |  |  |  |  |  |  |  |  |
| N | 100 | 100 | / |  | 70 | 72 | 58 | *χ*<sup>2</sup> | .423 |
| Age | 29.8<br>(±8.4) | 33.2<br>(±11.1) | *T* | **.015** | 32.7<br>(±11.7) | 29.8<br>(±9.1) | 32.1<br>(±8.5) | *F* | .203 |
| Gender (♀) | 42.0% | 40.0% | *χ*<sup>2</sup> | .886 | 42.9% | 40.3% | 39.7% | *χ*<sup>2</sup> | .924 |
| Education |  |  | *χ*<sup>2</sup> | .214 |  |  |  | *χ*<sup>2</sup> | .184 |
| Ongoing | 1.0% | 0.0% |  |  | 0.0% | 1.4% | 0.0% |  |  |
| Low | 2.0% | 2.0% |  |  | 2.9% | 0.0% | 3.4% |  |  |
| Middle | 11.0% | 21.0% |  |  | 20.0% | 8.3% | 20.7% |  |  |
| High | 78.0% | 70.0% |  |  | 71.4% | 84.7% | 63.8% |  |  |
| **Alcohol-Related Measures** |  |  |  |  |  |  |  |  |  |
| AUD Criteria | 0.1<br>(±0.2) | 4.2<br>(±1.8) | *T* | **<.001** | 2.0<br>(±2.5) | 2.2<br>(±2.4) | 2.1<br>(±2.6) | *F* | .892 |
| AUDIT | 4.0<br>(±3.0) | 14.9<br>(±4.9) | *T* | **<.001** | 8.8<br>(±7.6) | 10.0<br>(±6.4) | 9.5<br>(±6.2) | *F* | .612 |
| AUDIT-C | 2.8<br>(±1.5) | 6.8<br>(±1.7) | *T* | **<.001** | 4.4<br>(±2.9) | 5.0<br>(±2.4) | 5.2<br>(±2.4) | *F* | .225 |
| Consumption |  |  |  |  |  |  |  |  |  |
| grams/day | 32.7<br>(±23.5) | 75.9<br>(±40.4) | *T* | **<.001** | 48.6<br>(±42.1) | 55.3<br>(±38.1) | 60.3<br>(±37.7) | *F* | .250 |
| days/week | 1.1<br>(±1.2) | 4.1<br>(±1.8) | *T* | **<.001** | 2.5<br>(±2.3) | 2.7<br>(±2.0) | 2.8<br>(±2.1) | *F* | .717 |
| CAS-A | 3.0<br>(±5.3) | 16.2<br>(±9.7) | *T* | **<.001** | 9.3<br>(±10.6) | 9.4<br>(±10.0) | 10.2<br>(±10.2) | *F* | .886 |
| **Smoking-Related Measures** |  |  |  |  |  |  |  |  |  |
| TUD Criteria | 1.8<br>(±2.3) | 1.8<br>(±2.3) | *T* | .925 | 0.0<br>(±0.3) | 1.7<br>(±1.9) | 4.0<br>(±2.2) | *F* | **<.001** |
| FTND | 1.2<br>(±2.1) | 1.0<br>(±2.0) | *T* | .620 | 0.1<br>(±0.5) | 0.2<br>(±0.6) | 3.3<br>(±2.6) | *F* | **<.001** |
| Consumption |  |  |  |  |  |  |  |  |  |
| units/day | 4.8<br>(±6.3) | 5.2<br>(±7.2) | *T* | .675 | 0.0<br>(±0.0) | 3.0<br>(±2.7) | 13.3<br>(±6.7) | *F* | **<.001** |
| days/week | 2.5<br>(±3.1) | 2.7<br>(±3.1) | *T* | .796 | 0.0<br>(±0.0) | 1.6<br>(±1.9) | 7.0<br>(±0.0) | *F* | **<.001** |
| QSU-G | 2.6<br>(±1.3) | 2.9<br>(±1.3) | *T* | .313 | 2.8<br>(±0.0) | 1.9<br>(±1.0) | 3.5<br>(±1.2) | *F* | **<.001** |

*Note.* AUD = Alcohol Use Disorder; AUDIT = AUD Identification Test; AUDIT-C = AUDIT-Consumption; CAS-A = Craving Automated Scale for Alcohol; TUD = Tobacco Use Disorder; QSU-G = Questionnaire on Smoking Urges - German.

By design, the AUD group met more AUD criteria and showed higher AUDIT total scores than the non-AUD group (DSM-5 criteria: 4.2 vs. 0.1; AUDIT: 14.9 vs. 4.0), with most AUD participants (76%) falling within the mild-to-moderate severity range (2-5 DSM-5 criteria; see Appendix S6). Likewise, the number of TUD criteria and the FTND scores decreased from daily smokers to occasional smokers to non-smokers (DSM-5 criteria: 4.0 vs. 1.7 vs. 0.0; FTND: 3.3 vs. 0.2 vs. 0.1). In line with the orthogonal stratification design, the AUD groups did not differ in the number of TUD criteria or FTND scores, nor did the smoking status groups differ in AUD criteria or AUDIT scores. For a more detailed description of these groups and an overview across all six stratification groups, see Table 1 and S6.1, respectively.

### Behavioral Results

As summarized in the Supplementary Material S4, the initial task manipulation check revealed a significant PIT interference effect across the entire sample, *V* = 10384, *p* < .001, *r* = .56, reflected by a significant increase in the mean ER from the congruent (*M* = 16.46%, *SD* = 14.03%) to the incongruent condition (*M* = 30.74%, *SD* = 27.51%).

#### Associations With AUD and Smoking Status

As expected, the magnitude of this PIT interference effect (Δ*ER*) was significantly stronger in individuals with AUD (*M* = 17.65%, *SD* = 29.77%) than in individuals without AUD (*M* = 10.90%, *SD* = 24.80%), *β* = .03, *SE* = .01, *p*(one-sided) = .017, 95% CI [.002, .07] (see Figure 3A-B). In contrast, PIT interference did not increase significantly with increasing smoking status (Figure 3D-E), *β* = .001, *SE* = .01, *p*(one-sided) = .469, 95% CI [−.02, .02], nor did we observe a significant interaction between AUD and smoking status (see Figure S7.1), *β* = −.02, *SE* = .02, *p*(two-sided) = .242, 95% CI [−.06, .02]. An age- and gender-adjusted model further provided no evidence for significant age, *β* = .001, *SE* = .001, *p*(two-sided) = .351, 95% CI [−.001, .004], or gender effects, *β* = .01, *SE* = .01, *p*(two-sided) = .587, 95% CI [−.03, .04], on PIT interference. Notably, the AUD coefficient remained significant after adjusting for age and gender, *β* = .03, *SE* = .01, *p*(one-sided) = .038, 95% CI [−.002, .06].

**Figure 3.**
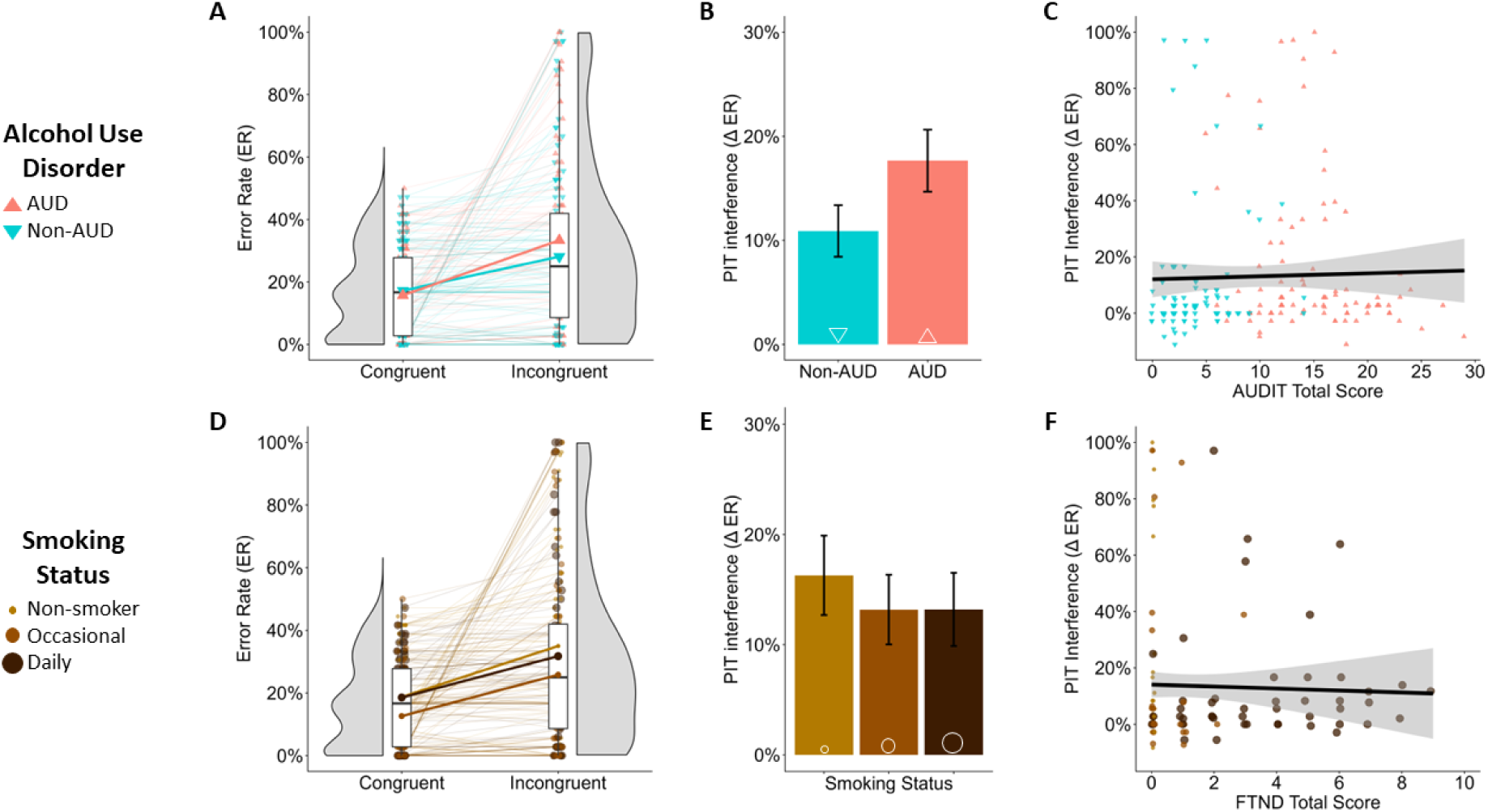
Association of PIT Interference With AUD, Smoking Status, and Disorder Severity. *Note.* Panel A: Distribution of participants’ mean error rates (ER) in the congruent and incongruent conditions, shown as half-violin plots and boxplots. Small colored triangles represent participants’ individual ERs, with thin lines illustrating the change in ER from the congruent to the incongruent condition. Larger triangles connected by solid lines indicate the corresponding group means and changes for the AUD and non-AUD groups. Panel B: Mean PIT interference effects (Δ*ER* = *ER_incongruent_ − ER_congruent_*) and standard errors for the AUD and Non-AUD groups. Panel C: Association between AUD severity (i.e., AUDIT total score) and PIT interference. Panel D: Distribution of participants’ mean error rates (ER) in the congruent and incongruent conditions, shown as half-violin plots and boxplots. Small colored points represent participants’ individual ERs, with thin lines illustrating the change in ER from the congruent to the incongruent condition. Larger colored points, connected by solid lines, indicate the corresponding group means and changes for each smoking-status group. Panel E: Mean PIT interference effects (Δ*ER* = *ER_incongruent_ − ER_congruent_*) and standard errors for the three smoking-status groups. Panel C: Association between TUD severity (i.e., FTND total score) and PIT interference.

#### Association With Continuous Substance-Related Measures

With respect to the hypothesized dimensional associations, summarized in Table 2, we found significant positive correlations between higher PIT interference and higher AUD severity, higher alcohol craving, and higher alcohol consumption, particularly with drinking frequency (see Figure 3C and 4A). Conversely, PIT interference did not correlate significantly with higher drinking quantity, nor did it predict drinking trajectories beyond baseline, *β* = .08, *SE* = .13, *p*(two-sided) = .576, 95% CI [−.23, .38], with overall drinking decreasing significantly across time points, *β* = −.15, *SE* = .04, *p*(two-sided) < .001, 95% CI [−.23, −.07].

**Figure 4.**
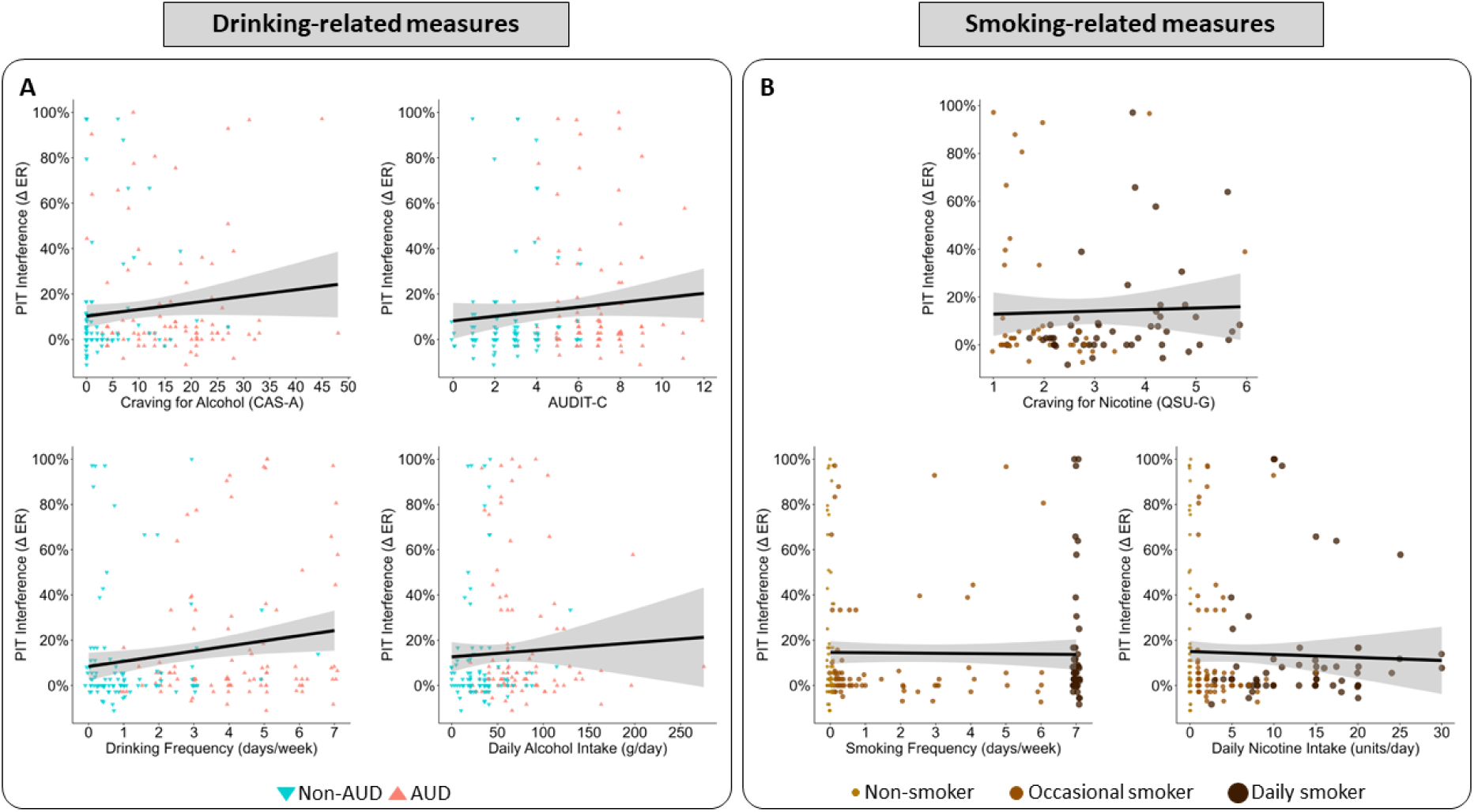
Association of PIT Interference With Substance-Specific Craving and Consumption Levels. Note. Each colored triangle or dot represents an individual participant, with solid lines indicating the association between PIT interference and continuous substance-related measures across the entire sample. Panel A: Stronger PIT interference was associated with higher levels of self-reported craving for alcohol (top left), higher levels of self-reported drinking (AUDIT-C; top right), more frequent drinking (bottom left), and higher quantities of alcohol intake per drinking day (bottom right). Panel B: Stronger PIT interference was not significantly associated with higher levels of self-reported craving for nicotine (QSU-G; top), more frequent smoking (bottom left), or higher amounts of tobacco products consumed per day (bottom right)

**Table 2.** Associations Between Behavioral PIT Interference and Continuous Substance-Related Measures.

|  | *τ* | *p* |
|---|---|---|
| **Continuous Alcohol-Related Measures** |  |  |
| AUD Severity (AUDIT) | .09 | **.044** |
| Drinking Frequency (days/week) | .12 | **.010** |
| Drinking Quantity (grams/day) | .08 | .057 |
| Overall Consumption (AUDIT-C) | .11 | **.022** |
| Alcohol Craving (CAS-A) | .11 | **.016** |
| **Continuous Smoking-Related Measures** |  |  |
| TUD severity (FTND) | .05 | .186 |
| Smoking Frequency (days/week) | .01 | .392 |
| Smoking Quantity (units/day) | .01 | .390 |
| Nicotine Craving (QSU-G) | .06 | .222 |

*Note.* Kendall’s Tau (*τ*) correlations were tested one-sided according to a priori directional hypotheses. AUD = Alcohol Use Disorder; AUDIT = AUD Identification Test; AUDIT-C = AUDIT-Consumption; CAS-A = Craving Automated Scale for Alcohol; FTND = Fagerström Test for Nicotine Dependence; g = grams; QSU-G = Questionnaire on Smoking Urges - German.

For smoking, we found no evidence for a significant association between greater PIT interference and higher TUD severity, smoking frequency, smoking quantity, or nicotine craving (see Figure 3F and 4B).

### Imaging Results

#### Neural PIT Interference

##### ROI Analysis

At the neural level, we found that incongruent PIT trials were associated with stronger brain responses in the right lPFC, *t*(179) = 2.58, *p*_FDR_(one-sided) = .009, Cohen’s *d* = 0.19, bilateral dmPFC, *t*(179) = 2.40, *p*_FDR_(one-sided) = .009, Cohen’s *d* = 0.18, and bilateral ACC, *t*(179) = 4.42, *p*(one-sided) < .001, Cohen’s *d* = 0.33, than congruent PIT trials. Unexpectedly, however, neural PIT interference did not correlate significantly with behavioral PIT interference in any of the included ROIs (*p* > .401; see Table S8.1). There was also no evidence for the hypothesized AUD group differences within the right lPFC, *t*(171.27) = 1.14, *p*_FDR_(one-sided) = .348, Cohen’s *d* = 0.17, bilateral dmPFC, *t*(173.41) = 0.94, *p*_FDR_(one-sided) = .348, Cohen’s *d* = 0.14, bilateral amygdala, *t*(177.17) = 0.91, *p*_FDR_(one-sided) = .817, Cohen’s *d* = 0.14, bilateral VS, *t*(174.75) = 0.67, *p*_FDR_(one-sided) = .817, Cohen’s *d* = 0.10, or the exploratory ACC mask, *t*(170.88) = 0.51, *p*(one-sided) = .307, Cohen’s *d* = 0.08. Likewise, none of these regions showed a significant association between neural PIT interference and smoking status (lPFC, dmPFC, VS, amygdala: *p*_FDR_ *≥* .465; ACC: *β* = −.04, *SE* = .14, *p*(one-sided) = .395, 95% CI [−0.31, 0.24]) or a significant AUD x smoking status interaction (lPFC, dmPFC, VS, amygdala: *p*_FDR_ = .981; ACC: *β* = .09, *SE* = .20, *p*(two-sided) = .670, 95% CI [−0.30, 0.48]; see Supplementary Material S8).

##### Whole Brain Analysis

As illustrated in Figure 5, the exploratory whole-brain analysis of neural PIT interference, with a significance threshold of *p* < .001 and cluster threshold of *k* > 50, largely converged with the ROI results, revealing significantly higher activation in the lPFC, dmPFC, and ACC during incongruent than congruent PIT trials. Beyond these regions, greater activation during incongruent vs. congruent PIT trials was also found in the right frontal pole, the angular gyri, and the insular cortices (see Table S8.2). In contrast, no brain region exhibited greater activation during congruent than incongruent PIT trials.

**Figure 5.**
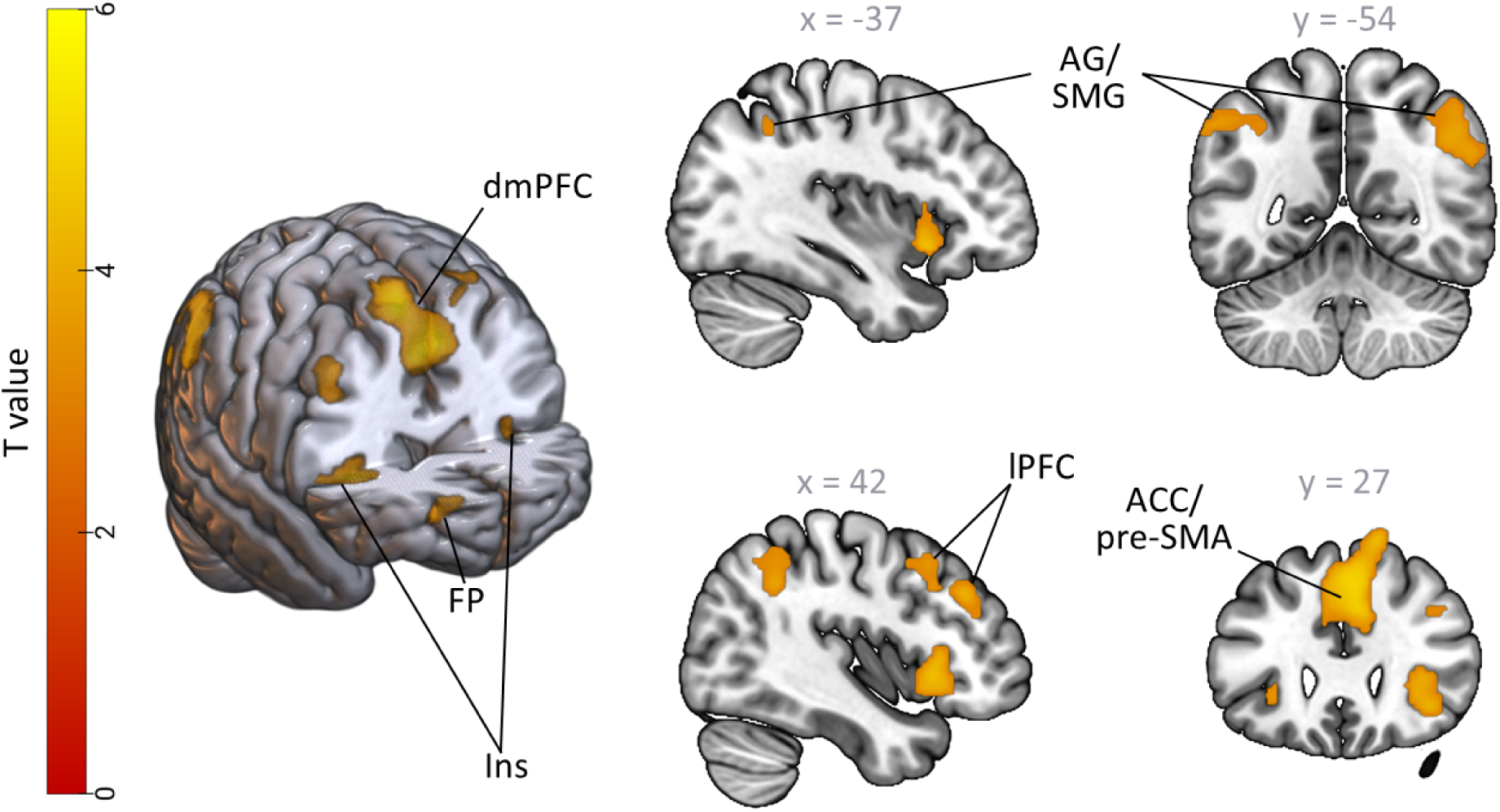
Exploratory Whole-Brain Analysis of Neural PIT Interference. *Note.* Relative to congruent PIT trials, incongruent PIT trials elicited greater activation within a frontoparietal network encompassing the right lateral prefrontal cortex (lPFC), the anterior cingulate cortex (ACC)/pre-supplementary motor area (pre-SMA), extending into the dorsomedial prefrontal cortex (dmPFC), the frontal pole (FP), the angular (AG) and supramarginal gyri (SMG). A significant neural PIT interference effect was also observed with the bilateral insular cortices (Ins). No brain regions showed greater activation during congruent than incongruent PIT trials.

#### Neural Incongruity

##### Association With AUD and Smoking Status

With respect to the more reliable incongruity contrast [40], previously used by PIT studies in risky drinkers [13, 14], individuals with AUD, relative to those without, exhibited significantly reduced activations within the right lPFC, *t*(177.31) = 2.16, *p*_FDR_(one-sided) = .032 *d* = 0.32, and bilateral dmPFC, *t*(177.97) = 2.16 *p*_FDR_(one-sided) = .032, *d* = 0.32, when exposed to a conflict between Pavlovian CS and instrumental control (see Figure 6A). The same group difference was also observed within the exploratory ACC ROI, *t*(174.78) = 2.44 *p*(one-sided) = .008, *d* = 0.36. However, individuals with AUD did not show significantly higher activation in the VS, *t*(177.92) = 0.02, *p*_FDR_(one-sided) = .508, *d* = 0.003, or amygdala, *t*(177.64) = −.57, *p*_FDR_(one-sided) = .377, *d* = −0.09, during incongruent PIT trials. Contrary to our hypothesis, there was also no evidence for an association of neural incongruity with increasing smoking status within any of pre-selected ROIs (lPFC: *β* = −.35, *SE* = .33, *p*_FDR_(one-sided) = .289, 95% CI [−1.00, 0.30]; dmPFC: *β* = *−*0.72, *SE* = .39, *p*_FDR_(one-sided) = .132, 95% CI [−1.49, 0.03]; VS: *β* = −.37, *SE* = .27, *p*_FDR_(one-sided) = .920, 95% CI [−0.89, 0.13]; amygdala: *β* = −.52, *SE* = .42, *p*_FDR_(one-sided) = .920, 95% CI [−1.30, 0.32]). The interaction between AUD and smoking status was also non-significant for all ROIs (dlPFC: *β* = .60, *SE* = .47, *p*_FDR_(one-sided) = .275, 95% CI [−0.33, 1.51]; dmPFC: *β* = 1.35, *SE* = .55, *p*_FDR_(one-sided) = .061, 95% CI [0.30, 2.44]; VS: *β* = .60, *SE* = .38, *p*_FDR_(one-sided) = .233, 95% CI [−0.14, 1.31]; amygdala: *β* = .33, *SE* = .59, *p*_FDR_(one-sided) = .582, 95% CI [−0.83, 1.52]). Only without FDR correction, there was a significant association between smoking status and decreasing dmPFC activation during incongruent PIT trials, *p*(one-sided) = .033, and a significant interaction between AUD and smoking status within the dmPFC, *p*(two-sided) = .015. A similar pattern of decreasing neural activation during incongruent PIT trials with increasing smoking status was also observed within the ACC, *β* = −.65, *SE* = .33, *p*(one-sided) = .023, 95% CI [−1.35, −0.04]. However, the AUD x smoking status interaction in the ACC did not reach significance, *β* = .89, *SE* = .46, *p*(two-sided) = .058, 95% CI [−0.02, 1.78].

**Figure 6.**
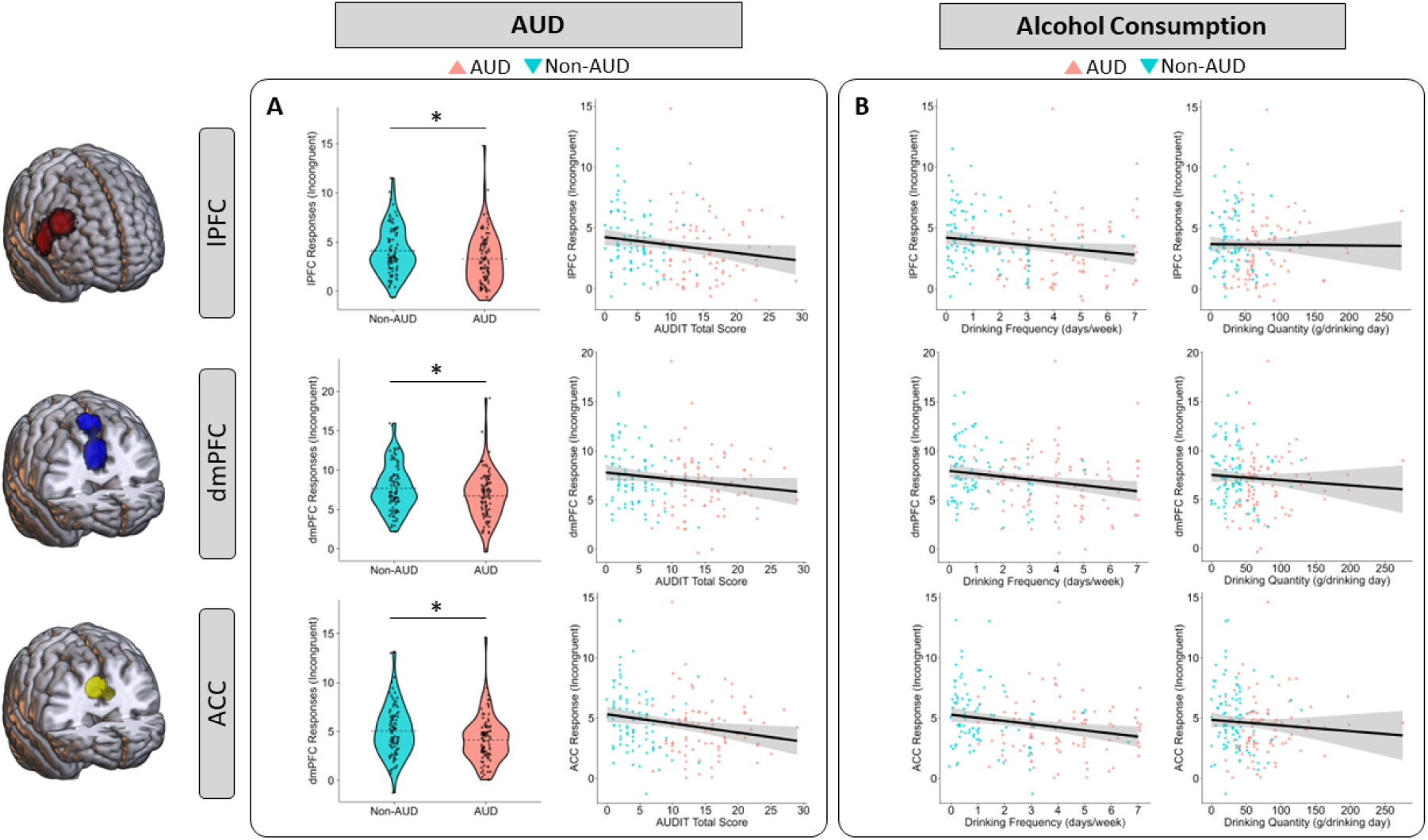
Association of Prefrontal Neural Activation During Incongruent PIT Trials with AUD, AUD severity, and Alcohol Consumption. *Note.* Each colored triangle or dot represents an individual participant, with solid lines indicating the associations between incongruity-related activation, AUD, and alcohol consumption levels. Panel A: Individuals with AUD, relative to those without, exhibited significantly reduced activation during incongruent PIT trials within the right lPFC (top left), bilateral dmPFC (middle left), and bilateral ACC (bottom left), with higher levels of AUD severity correlating negatively with incongruity-related activation in the lPFC (top right), dmPFC (middle right), and ACC (bottom right) across the entire sample (only without FDR-correction). Panel B: More frequent drinking, but not higher amounts of daily alcohol intake, were significantly associated with weaker brain responses within the right lPFC (top left), bilateral dmPFC (middle left), and bilateral ACC (bottom left) during incongruent PIT trials.

##### Association With Continuous Substance-Related Measures

At the dimensional level, only drinking frequency was significantly associated with neural incongruity after FDR correction (see Table 3. As illustrated in Figure 6B, higher drinking frequency correlated with weaker lPFC, dmPFC, and ACC activation, *r* = −.21, *p*(one-sided) = .002, during incongruent PIT trials.

**Table 3.** Association Between Neural Activation During Incongruent PIT Trials and Continuous Substance-Related Measures.

|  | **Alcohol** |  |  | **Smoking** |  |  |
|---|---|---|---|---|---|---|
|  | *r* | *p* | *p*<sub>FDR</sub> | *r* | *p* | *p*<sub>FDR</sub> |
| **Disorder Severity** | *AUDIT total* |  |  | *FTND total* |  |  |
| right lPFC | -.17 | **.014** | .056 | .001 | .504 | .975 |
| bilateral dmPFC | -.15 | **.028** | .056 | -.11 | .082 | .329 |
| bilateral VS | .06 | .208 | .208 | -.11 | .907 | .975 |
| bilateral amygdala | .07 | .197 | .208 | -.16 | .975 | .975 |
| **Consumption Frequency** | *drinking days/week* |  |  | *smoking days/week* |  |  |
| right lPFC | -.16 | **.014** | **.027** | .01 | .563 | .763 |
| bilateral dmPFC | -.20 | **.003** | **.012** | -.03 | .350 | .763 |
| bilateral VS | -.02 | .597 | .597 | -.01 | .572 | .763 |
| bilateral amygdala | .01 | .469 | .597 | -.10 | .917 | .917 |
| **Consumption Quantity** | *g/day* |  |  | *units/day* |  |  |
| right lPFC | -.01 | .454 | .454 | -.04 | .307 | .614 |
| bilateral dmPFC | -.07 | .169 | .338 | -.06 | .207 | .614 |
| bilateral VS | .04 | .282 | .376 | -.05 | .726 | .968 |
| bilateral amygdala | .12 | **.049** | .195 | -.16 | .986 | .986 |
| **Craving** | *CAS-A sum* |  |  | *QSU-G mean* |  |  |
| right lPFC | -.10 | .100 | .200 | .04 | .626 | .834 |
| bilateral dmPFC | -.11 | .076 | .200 | -.12 | .130 | .520 |
| bilateral VS | .05 | .279 | .373 | .04 | .370 | .741 |
| bilateral amygdala | .02 | .390 | .390 | -.19 | .958 | .958 |

*Note. p*<sub>FDR</sub> = *p*-values corrected for multiple comparisons using the false discovery rate (FDR) across the four regions of interest (ROIs), separately for each substance-related variable. lPFC = lateral prefrontal cortex; dmPFC = dorsomedial prefrontal cortex; VS = ventral striatum.

An exploratory analysis without FDR correction further revealed a significant negative correlation between greater AUD severity and weaker lPFC and dmPFC (see Table 3), as well as ACC activation (*r* = −.20, *p*(one-sided) = .005) during incongruent PIT trials (see Figure 6A). For the ACC, significant negative correlations were also found with alcohol craving, *r* = −.16, *p*(one-sided) = .021, overall drinking (AUDIT-C), *r* = −.16, *p*(one-sided) = .021, and drinking frequency, *r* = −.21, *p*(one-sided) = .002. Without FDR correction, we also found a significant positive correlation between greater amygdala activation and drinking quantity.

In contrast, the correlations between incongruity-related ACC activation and drinking quantity, *r* = −.08, *p*(one-sided) = .156, nicotine craving, *r* = −.02, *p*(one-sided) = .436, smoking frequency, *r* = −.05, *p*(one-sided) = .249, smoking quantity, *r* = −.08, *p*(one-sided) = .143, and TUD severity (FTND), *r* = −.11, *p*(one-sided) = .095, did not reach significance.

## Discussion

In this study, we investigated whether enhanced PIT interference and its neural correlates – previously observed in risky drinkers and alcohol-dependent inpatients – generalize to non-treatment-seeking individuals with predominantly mild-to-moderate AUD, and whether enhanced PIT also extends to smoking. PIT interference was assessed in a general-population sample, which was stratified by AUD and smoking status during recruitment, using a joystick-based single-lever PIT paradigm. We found that participants with AUD exhibit greater PIT interference than participants without AUD, indicating an increased susceptibility to the influence of Pavlovian cues when they conflict with the required instrumental response. Greater PIT interference also correlated with higher AUD severity, alcohol craving, and drinking frequency. At the neural level, AUD was associated with weaker brain responses to PIT conflicts, particularly in the lPFC, dmPFC, and ACC, which correlated with higher drinking frequency. In contrast, we found no evidence for an association between greater PIT interference or related neural activation with smoking status, TUD severity, or other smoking-related measures.

This study thus replicates previous findings of enhanced PIT interference in AUD [12–14] and extends this literature to a broader, non-treatment-seeking AUD population that often goes clinically undetected [23, 24]. Together with the evidence from risky drinkers and alcohol-dependent inpatients [12, 13], our study suggests that an increased susceptibility to the interference between Pavlovian cues and instrumental control generalizes across the AUD spectrum, and that its magnitude increases with disorder severity. In everyday life, this reduced ability to shield goal-directed behavior from Pavlovian cues may lead to more frequent maladaptive approaches to harmful stimuli, including alcohol, and may thereby contribute to the development and maintenance of AUD. In line with this notion, Sommer et al. [12] and Chen et al. [14] observed an association between greater PIT interference and increases in alcohol consumption and relapse after detoxification. The present study adds to this literature by demonstrating a link between greater PIT interference and higher levels of alcohol craving and alcohol consumption, particularly drinking frequency, while controlling for smoking status.

Importantly, however, we found no evidence for an association of PIT interference with smoking status or other smoking-related measures. Based on the general nature of the associative learning mechanisms involved in PIT, it has been hypothesized that the relevance of enhanced PIT interference for drug use may extend to all substances, including tobacco [6, 7, 21, 25]. However, a previous study investigating the effects of smoking-related Pavlovian CSs on instrumental behavior indicated that the magnitude of the observed PIT effects may not differ between groups of varying smoking status [30]. The present study complements this finding by demonstrating that occasional and daily smokers are not more prone to inadequate approach-avoidance behavior than non-smokers when non-drug-related Pavlovian CSs conflict with the required instrumental response. Notably, the magnitude of PIT interference was also independent of TUD severity, smoking-related craving, and tobacco consumption levels. Taken together, PIT studies in human smokers thus challenge the hypothesis that greater PIT interference extends to smoking.

In addition to this dissociation at the behavioral level, AUD- and smoking status-related effects also diverged at the neural level, particularly during PIT conflicts. While the smoking status groups showed no significant differences in neural activation across all ROIs, participants with AUD exhibited weaker brain responses in the lPFC, dmPFC, and ACC during incongruent PIT trials than non-AUD participants. Weaker lPFC, dmPFC, and ACC activation also correlated with higher drinking frequency, while being independent of smoking-related measures. These findings are consistent with prior literature documenting differences in lPFC and dmPFC activation during PIT conflicts between high- and low-risk drinkers [13]. More importantly, they extend these findings to individuals with predominantly mild-to-moderate AUD, link these alterations to clinically relevant outcome measures, and implicate the ACC as a more anatomically circumscribed region within the larger dmPFC. As suggested by Chen et al. [13], these alterations may reflect dysregulation within a broader fronto-striatal-limbic network that integrates bottom-up motivational input, detects potential conflicts, and recruits top-down control resources if necessary to guide motor execution [57, 58]. Within this network, the lPFC, dmPFC, and ACC have consistently been linked to cognitive control functions, particularly performance monitoring, conflict detection (dmPFC, especially ACC), and response inhibition (lPFC) [59–61].

One possible explanation for the observed dissociation between AUD- and TUD-related effects is that cognitive control deficits may differ between both disorders. Numerous studies observed that individuals with AUD perform worse than individuals without AUD on tasks requiring the delay, withholding, or interruption of prepotent responses [62]. Other work demonstrated that individuals with AUD exhibit significant impairments in resolving conflicts between competing stimuli or response tendencies, a subfacet of inhibitory control known as interference control [63]. Given the co-occurrence of competing stimuli, the simultaneous activation of competing response tendencies, and the need to inhibit one of them during PIT conflicts, these AUD-related deficits in inhibitory control may be detrimental for resolving PIT interference [12, 13]. Unlike in AUD, the evidence for inhibitory control deficits in TUD and chronic smokers is considerably mixed [64–70]. Some studies even suggest that impairments in inhibitory control may emerge only after several hours of abstinence and when cognitive demands are low [71–73]. It is therefore possible that smokers in this study showed no inhibitory control impairments beyond those related to AUD. This interpretation aligns with the observed alterations in inhibitory-control-related brain regions in AUD, whose activation was independent of smoking status. It may also explain why including smoking status and a potential interaction between AUD and smoking did not alter the results for an AUD-related difference in PIT interference.

## Limitations

However, several limitations must be acknowledged.

First, there was no direct readout for the strength of the Pavlovian cue-outcome associations, leaving it unclear whether low ERs during PIT conflicts reflect good top-down control or insufficient Pavlovian learning. Similarly, it cannot be ruled out that maximal PIT interference may result from factors other than PIT conflicts, such as uncertainty about shell valence or misconceptions about its relationship with Pavlovian CSs.

Second, technical constraints of the MRI joystick introduced noise into data recordings, necessitating the exclusion of individual participants and trials.

Third, there was a small yet significant age difference between AUD groups that was not intended during recruitment.

Fourth, there was high within-group variability in smoking and TUD severity in the occasional smoker group, which may have limited our ability to detect smoking-status-related differences in PIT interference.

## Future Outlook & Implications

Building on the existing work in AUD, future studies should examine the temporal stability of PIT interference and assess how fluctuations across multiple PIT assessments may contribute to substance use, the development of an AUD, and relapse. In this context, they should also explore the role of inhibitory control deficits, specifically which subfacets of inhibitory control may be most relevant for overcoming PIT interference, and whether AUD-related impairments reflect state- or trait-like deficits that contribute to enhanced PIT. Similar to this study, future studies may also investigate whether an increased susceptibility to PIT and its neural correlates extend to the chronic use of substances with more severe and consistent negative effects on inhibitory control than smoking, such as cocaine [62]. Together, these studies may inform clinicians about whether they should focus on managing transient periods of heightened PIT or preventing the translation of a more stable vulnerability into drinking and, ultimately, AUD.

Despite these open questions, this study reaffirms the importance of considering PIT in the treatment of AUD. Recent evidence suggests that the impact of Pavlovian CSs may be reduced by cognitive bias modification interventions, thereby representing a promising adjunctive treatment approach for AUD [74]. So far, however, the evidence that these effects translate into sustained clinical outcomes, including long-term reductions in alcohol use, remains inconsistent [75, 76]. A more comprehensive understanding of the neurocognitive mechanisms underlying PIT may facilitate the refinement of such interventions and may eventually improve their effectiveness and the durability of their effects [74].

## Conclusion

Overall, this study adds to a growing literature implicating an increased impact of Pavlovian CSs on instrumental approach-avoidance behavior in AUD. It further underscores the clinical relevance of such effects by linking stronger PIT interference to higher alcohol craving, more frequent drinking, and greater AUD severity. Consistent with previous work, the neuroimaging results point to an impaired recruitment of prefrontal control resources during PIT conflicts as a potential neurocognitive mechanism underlying PIT. Most importantly, however, the results of this study challenge the hypothesis that enhanced PIT interference and related neural alterations extend to smoking.

## Acknowledgments

This research was funded by the German Research Foundation (Deutsche Forschungsgemeinschaft, DFG project numbers 178833530 [SFB 940], 402170461 [TRR 265], 454245598 [IRTG 2773], 460922097 [INST 269/881-1 FUGG], and 521379614 [TRR 393]).

We thank Clarissa Grundmann, Angela Hentschel, Julia Göstl, and Michaela Reuter for their support in participant recruitment and data collection.

## References

[1] World Health Organization. Global status report on alcohol and health and treatment of substance use disorders. 2024. url: https://iris.who.int/bitstream/handle/10665/377960/9789240096745-eng.pdf?sequence=1.

[2] Olderbak, S., Hollweck, R., Krowartz, E.-M., Möckl, J. and Hoch, E. ‘Psychoactive Substance Use in Germany: Findings From the Epidemiological Survey of Substance Abuse (ESA) in 2024’. In: Deutsches Ärzteblatt International 122 (2025), pp. 625–631. doi: 10.3238/arztebl.m2025.0157.

[3] Rehm, J., Shield, K. D., Rehm, M. X., Gmel, G. and Frick, U. Alcohol consumption, alcohol dependence, and attributable burden of disease in Europe: potential gains from effective interventions for alcohol dependence. CAMH, 2012.

[4] Mannes, Z. L., Shmulewitz, D., Livne, O., Stohl, M. and Hasin, D. S. ‘Correlates of mild, moderate, and severe Alcohol Use Disorder among adults with problem substance use: Validity implications for DSM-5’. In: Alcoholism: Clinical and Experimental Research 45.10 (2021), pp. 2118–2129. doi: 10.1111/acer.14701. eprint: https://onlinelibrary.wiley.com/doi/pdf/10.1111/acer.14701. url: https://onlinelibrary.wiley.com/doi/abs/10.1111/acer.14701.

[5] MacKillop, J., Agabio, R., Feldstein Ewing, S. W., Heilig, M., Kelly, J. F., Leggio, L. et al. ‘Hazardous drinking and alcohol use disorders’. In: Nature Reviews Disease Primers 8.80 (2022), pp. 1–25. doi: 10.1038/s41572-022-00406-1.

[6] Heinz, A., Beck, A., Halil, M. G., Pilhatsch, M., Smolka, M. N. and Liu, S. ‘Addiction as Learned Behavior Patterns’. In: Journal of Clinical Medicine 8.8, 1086 (2019). doi: 10.3390/jcm8081086.

[7] Robinson, T. E. and Berridge, K. C. ‘The neural basis of drug craving: An incentive-sensitization theory of addiction’. In: Brain Research Reviews 18.3 (1993), pp. 247–291. issn: 0165-0173. doi: 10.1016/0165-0173(93)90013-P. url: https://www.sciencedirect.com/science/article/pii/016501739390013P.

[8] Cartoni, E., Balleine, B. and Baldassarre, G. ‘Appetitive Pavlovian-instrumental Transfer: A review’. In: Neuroscience & Biobehavioral Reviews 71 (2016), pp. 829–848. issn: 0149-7634. doi: 10.1016/j.neubiorev.2016.09.020. url: https://www.sciencedirect.com/science/article/pii/S0149763416300616.

[9] Garbusow, M., Schad, D. J., Sommer, C., Jünger, E., Sebold, M., Friedel, E. et al. ‘Pavlovian-to-Instrumental Transfer in Alcohol Dependence: A Pilot Study’. In: Neuropsychobiology 70.2 (Oct. 2014), pp. 111–121. issn: 0302-282X. doi: 10.1159/000363507. eprint: https://karger.com/nps/article-pdf/70/2/111/3259133/000363507.pdf. url: 10.1159/000363507.

[10] Garbusow, M., Schad, D. J., Sebold, M., Friedel, E., Bernhardt, N., Koch, S. P. et al. ‘Pavlovian-to-instrumental transfer effects in the nucleus accumbens relate to relapse in alcohol dependence’. In: Addiction Biology 21.3 (2016), pp. 719–731. doi: 10.1111/adb.12243. eprint: https://onlinelibrary.wiley.com/doi/pdf/10.1111/adb.12243. url: https://onlinelibrary.wiley.com/doi/abs/10.1111/adb.12243.

[11] Chen, K., Schlagenhauf, F., Sebold, M., Kuitunen-Paul, S., Chen, H., Huys, Q. J. et al. ‘The Association of Non–Drug-Related Pavlovian-to-Instrumental Transfer Effect in Nucleus Accumbens With Relapse in Alcohol Dependence: A Replication’. In: Biological Psychiatry 93.6 (2023). Addiction Phenotypes and Mechanisms, pp. 558–565. issn: 0006-3223. doi: 10.1016/j.biopsych.2022.09.017. url: https://www.sciencedirect.com/science/article/pii/S0006322322015992.

[12] Sommer, C., Garbusow, M., Juenger, E., Pooseh, S., Bernhardt, N., Birkenstock, J. et al. ‘Strong seduction: impulsivity and the impact of contextual cues on instrumental behavior in alcohol dependence’. In: Translational Psychiatry 7, e1183 (2017). doi: 10.1038/tp.2017.158.

[13] Chen, H., Nebe, S., Mojtahedzadeh, N., Kuitunen-Paul, S., Garbusow, M., Schad, D. J. et al. ‘Susceptibility to interference between Pavlovian and instrumental control is associated with early hazardous alcohol use’. In: Addiction Biology 26.4, e12983 (2021). doi: 10.1111/adb.12983. eprint: https://onlinelibrary.wiley.com/doi/pdf/10.1111/adb.12983. url: https://onlinelibrary.wiley.com/doi/abs/10.1111/adb.12983.

[14] Chen, H., Belanger, M. J., Garbusow, M., Kuitunen-Paul, S., Huys, Q. J. M., Heinz, A. et al. ‘Susceptibility to interference between Pavlovian and instrumental control predisposes risky alcohol use developmental trajectory from ages 18 to 24’. In: Addiction Biology 28.2, e13263 (2023). doi: 10.1111/adb.13263. eprint: https://onlinelibrary.wiley.com/doi/pdf/10.1111/adb.13263. url: https://onlinelibrary.wiley.com/doi/abs/10.1111/adb.13263.

[15] Garbusow, M., Nebe, S., Sommer, C., Kuitunen-Paul, S., Sebold, M., Schad, D. J. et al. ‘Pavlovian-To-Instrumental Transfer and Alcohol Consumption in Young Male Social Drinkers: Behavioral, Neural and Polygenic Correlates’. In: Journal of Clinical Medicine 8.8, 1188 (2019). doi: 10.3390/jcm8081188.

[16] Hall, J., Parkinson, J. A., Connor, T. M., Dickinson, A. and Everitt, B. J. ‘Involvement of the central nucleus of the amygdala and nucleus accumbens core in mediating Pavlovian influences on instrumental behaviour’. In: European Journal of Neuroscience 13.10 (2001), pp. 1984–1992. doi: 10.1046/j.0953-816x.2001.01577.x. url: https://onlinelibrary.wiley.com/doi/abs/10.1046/j.0953-816x.2001.01577.x.

[17] Geurts, D. E. M., Huys, Q. J. M., Ouden, H. E. M. den and Cools, R. ‘Aversive Pavlovian Control of Instrumental Behavior in Humans’. In: Journal of Cognitive Neuroscience 25.9 (Sept. 2013), pp. 1428–1441. issn: 0898-929X. doi: 10.1162/jocn_a_00425. eprint: https://direct.mit.edu/jocn/article-pdf/25/9/1428/1945461/jocn_a_00425.pdf. url: 10.1162/jocn_a_00425.

[18] Mendelsohn, A., Pine, A. and Schiller, D. ‘Between thoughts and actions: motivationally salient cues invigorate mental action in the human brain’. In: Neuron 81.1 (2014), pp. 207–217. doi: 10.1016/j.neuron.2013.10.019.

[19] Prévost, C., Liljeholm, M., Tyszka, J. M. and O’Doherty, J. P. ‘Neural Correlates of Specific and General Pavlovian-to-Instrumental Transfer within Human Amygdalar Subregions: A High-Resolution fMRI Study’. In: Journal of Neuroscience 32.24 (2012), pp. 8383–8390. issn: 0270-6474. doi: 10.1523/JNEUROSCI.6237-11.2012. eprint: https://www.jneurosci.org/content/32/24/8383.full.pdf. url: https://www.jneurosci.org/content/32/24/8383.

[20] Talmi, D., Seymour, B., Dayan, P. and Dolan, R. J. ‘Human Pavlovian–Instrumental Transfer’. In: Journal of Neuroscience 28.2 (2008), pp. 360–368. issn: 0270-6474. doi: 10.1523/JNEUROSCI.4028-07.2008. eprint: https://www.jneurosci.org/content/28/2/360.full.pdf. url: https://www.jneurosci.org/content/28/2/360.

[21] Robinson, M. J. F., Fischer, A. M., Ahuja, A., Lesser, E. N. and Maniates, H. ‘Roles of “Wanting” and “Liking” in Motivating Behavior: Gambling, Food, and Drug Addictions’. In: Behavioral Neuroscience of Motivation. Ed. By Simpson, E. H. and Balsam, P. D. Cham: Springer, 2016, pp. 105–136. isbn: 978-3-319-26935-1. doi: 10.1007/7854_2015_387. url: 10.1007/7854_2015_387.

[22] Hung, Y., Gaillard, S. L., Yarmak, P. and Arsalidou, M. ‘Dissociations of cognitive inhibition, response inhibition, and emotional interference: Voxelwise ALE meta-analyses of fMRI studies’. In: Human Brain Mapping 39.10 (2018), pp. 4065–4082. doi: 10.1002/hbm.24232. eprint: https://onlinelibrary.wiley.com/doi/pdf/10.1002/hbm.24232. url: https://onlinelibrary.wiley.com/doi/abs/10.1002/hbm.24232.

[23] Grant, B. F., Goldstein, R. B., Saha, T. D., Chou, S. P., Jung, J., Zhang, H. et al. ‘Epidemiology of DSM-5 Alcohol Use Disorder: Results From the National Epidemiologic Survey on Alcohol and Related Conditions III’. In: JAMA Psychiatry 72.8 (2015), pp. 757–766. doi: 10.1001/jamapsychiatry.2015.0584.

[24] Probst, C., Manthey, J., Martinez, A. and Rehm, J. ‘Alcohol use disorder severity and reported reasons not to seek treatment: A cross-sectional study in European primary care practices’. In: Substance Abuse Treatment Prevention and Policy 10.32 (2015), pp. 1–10. doi: 10.1186/s13011-015-0028-z.

[25] Garbusow, M., Ebrahimi, C., Riemerschmid, C., Daldrup, L., Rothkirch, M., Chen, K. et al. ‘Pavlovian-to-Instrumental Transfer across Mental Disorders: A Review’. In: Neuropsychobiology 81.5 (July 2022), pp. 418–437. doi: 10.1159/000525579.

[26] Hogarth, L., Dickinson, A., Wright, A., Kouvaraki, M. and Duka, T. ‘The role of drug expectancy in the control of human drug seeking.’ In: Journal of Experimental Psychology: Animal Behavior Processes 33.4 (2007), pp. 484–496. doi: 10.1037/0097-7403.33.4.484.

[27] Hogarth, L. ‘Goal-Directed and Transfer-Cue-Elicited Drug-Seeking are Dissociated by Pharmacotherapy: Evidence for Independent Additive Controllers’. In: Journal of Experimental Psychology: Animal Behavior Processes 38.3 (2012), pp. 266–278. doi: 10.1037/a0028914.

[28] Hogarth, L., Maynard, O. M. and Munafò, M. R. ‘Plain cigarette packs do not exert Pavlovian to instrumental transfer of control over tobacco-seeking’. In: Addiction 110.1 (2015), pp. 174–182. doi: 10.1111/add.12756. eprint: https://onlinelibrary.wiley.com/doi/pdf/10.1111/add.12756. url: https://onlinelibrary.wiley.com/doi/abs/10.1111/add.12756.

[29] Hogarth, L., Field, M. and Rose, A. K. ‘Phasic transition from goal-directed to habitual control over drug-seeking produced by conflicting reinforcer expectancy’. In: Addiction Biology 18.1 (2013), pp. 88–97. doi: 10.1111/adb.12009. eprint: https://onlinelibrary.wiley.com/doi/pdf/10.1111/adb.12009. url: https://onlinelibrary.wiley.com/doi/abs/10.1111/adb.12009.

[30] Hogarth, L. and Chase, H. ‘Evaluating Psychological Markers for Human Nicotine Dependence: Tobacco Choice, Extinction, and Pavlovian-to-Instrumental Transfer’. In: Experimental and Clinical Psychopharmacology 20.3 (Feb. 2012), pp. 213–224. doi: 10.1037/a0027203.

[31] Manglani, H. R., Lewis, A. H., Wilson, S. J. and Delgado, M. R. ‘Pavlovian-to-Instrumental Transfer of Nicotine and Food Cues in Deprived Cigarette Smokers’. In: Nicotine & Tobacco Research 19.6 (May 2017), pp. 670–676. issn: 1462-2203. doi: 10.1093/ntr/ntx007. eprint: https://academic.oup.com/ntr/article-pdf/19/6/670/24365809/ntx007.pdf. url: 10.1093/ntr/ntx007.

[32] Falk, D. E., Yi, H.-y. and Hiller-Sturmhöfel, S. ‘An epidemiologic analysis of co-occurring alcohol and tobacco use and disorders: findings from the National Epidemiologic Survey on Alcohol and Related Conditions’. In: Alcohol Research & Health 29.3 (2006), pp. 162–171.

[33] American Psychiatric Association. Diagnostic and statistical manual of mental disorders. 5th ed. American Psychiatric Association, 2013. doi: 10.1176/appi.books.9780890425596.

[34] Kuitunen-Paul, S., Rehm, J., Lachenmeier, D. W., Kadrić, F., Kuitunen, P. T., Wittchen, H.-U. et al. ‘Assessment of alcoholic standard drinks using the Munich composite international diagnostic interview (M-CIDI): An evaluation and subsequent revision’. In: International Journal of Methods in Psychiatric Research 26.3, e1563 (2017). doi: 10.1002/mpr.1563.

[35] Saunders, J. B., Aasland, O. G., Babor, T. F., De La Fuente, J. R. and Grant, M. ‘Development of the Alcohol Use Disorders Identification Test (AUDIT): WHO Collaborative Project on Early Detection of Persons with Harmful Alcohol Consumption-II’. In: Addiction 88.6 (1993), pp. 791–804. doi: 10.1111/j.1360-0443.1993.tb02093.x. eprint: https://onlinelibrary.wiley.com/doi/pdf/10.1111/j.1360-0443.1993.tb02093.x. url: https://onlinelibrary.wiley.com/doi/abs/10.1111/j.1360-0443.1993.tb02093.x.

[36] Heatherton, T. F., Kozlowski, L. T., Frecker, R. C. and Fagerstrom, K.-O. ‘The Fagerström test for nicotine dependence: a revision of the Fagerstrom Tolerance Questionnaire’. In: British journal of addiction 86.9 (1991), pp. 1119–1127. doi: 10.1111/j.1360-0443.1991.tb01879.x.

[37] Vollstädt-Klein, S., Leménager, T., Jorde, A., Kiefer, F. and Nakovics, H. ‘Development and Validation of the Craving Automated Scale for Alcohol’. In: Alcoholism: Clinical and Experimental Research 39.2 (2015), pp. 333–342. doi: 10.1111/acer.12636. eprint: https://onlinelibrary.wiley.com/doi/pdf/10.1111/acer.12636. url: https://onlinelibrary.wiley.com/doi/abs/10.1111/acer.12636.

[38] Müller, V., Mucha, R. F., Ackermann, K. and Pauli, P. ‘Die Erfassung Des Cravings Bei Rauchern Mit Einer Deutschen Version Des “Questionnaire on Smoking Urges” (QSU-G)Die Erfassung Des Cravings Bei Rauchern Mit Einer Deutschen Version Des “Questionnaire on Smoking Urges” (QSU-G) [Assessment of craving in smokers using a German version of the Questionnaire on Smoking Urges (QSU-G)]’. In: Zeitschrift für Klinische Psychologie Und Psychotherapie 30.3 (2001), pp. 164–171. doi: 10.1026/0084-5345.30.3.164.

[39] Belanger, M. J., Chen, H., Hentschel, A., Garbusow, M., Ebrahimi, C., Knorr, F. G. et al. ‘Development of Novel Tasks to Assess Outcome-Specific and General Pavlovian-to-Instrumental Transfer in Humans’. In: Neuropsychobiology 81.5 (Nov. 2022), pp. 370–386. issn: 0302-282X. doi: 10.1159/000526774. eprint: https://karger.com/nps/article-pdf/81/5/370/3731083/000526774.pdf. url: 10.1159/000526774.

[40] Belanger, M. J., Chen, H., Fröhner, J. H., Garbusow, M., Heinz, A. and Smolka, M. N. ‘Behavioural and Neural Reliability of a Pavlovian-to-Instrumental Transfer Task’. In: Addiction Biology 30.12, e70112 (2025). e70112 9961735. doi: 10.1111/adb.70112. eprint: https://onlinelibrary.wiley.com/doi/pdf/10.1111/adb.70112. url: https://onlinelibrary.wiley.com/doi/abs/10.1111/adb.70112.

[41] Brainard, D. H. and Vision, S. ‘The psychophysics toolbox’. In: Spatial vision 10.4 (1997), pp. 433–436.

[42] Kleiner, M., Brainard, D., Pelli, D., Ingling, A., Murray, R. and Broussard, C. ‘What’s new in Psychtoolbox-3’. In: Perception 36 (Jan. 2007), pp. 1–16.

[43] Pelli, D. G. and Vision, S. ‘The VideoToolbox software for visual psychophysics: Transforming numbers into movies’. In: Spatial vision 10 (1997), pp. 437–442.

[44] Sliedrecht, W., de Waart, R., Witkiewitz, K. and Roozen, H. G. ‘Alcohol use disorder relapse factors: A systematic review’. In: Psychiatry Research 278 (2019), pp. 97–115. issn: 0165-1781. doi: 10.1016/j.psychres.2019.05.038. url: https://www.sciencedirect.com/science/article/pii/S0165178119303841.

[45] Martins, J. S., Fogelman, N., Wemm, S., Hwang, S. and Sinha, R. ‘Alcohol craving and withdrawal at treatment entry prospectively predict alcohol use outcomes during outpatient treatment’. In: Drug and Alcohol Dependence 231, 109253 (2022). issn: 0376-8716. doi: 10.1016/j.drugalcdep.2021.109253.

[46] Gass, J. C., Motschman, C. A. and Tiffany, S. T. ‘The relationship between craving and tobacco use behavior in laboratory studies: a meta-analysis.’ In: Psychology of Addictive Behaviors 28.4 (2014), pp. 1162–1176. doi: 10.1037/a0036879.

[47] Vafaie, N. and Kober, H. ‘Association of drug cues and craving with drug use and relapse: a systematic review and meta-analysis’. In: JAMA psychiatry 79.7 (2022), pp. 641–650. doi: 10.1001/jamapsychiatry.2022.1240.

[48] Venables, W. N. and Ripley, B. D. Modern Applied Statistics with S. 4th ed. New York: Springer, 2002. url: https://www.stats.ox.ac.uk/pub/MASS4/.

[49] Höfler, M. ‘Robust tests should be the default, not the backup’. en. In: Peer Community Journal 6, e1 (2026). doi: 10.24072/pcjournal.670. url: https://peercommunityjournal.org/articles/10.24072/pcjournal.670/.

[50] Leonhart, R. Lehrbuch Statistik: Einstieg und Vertiefung[Textbook of Statistics: Introduction and Deepening]. 4th ed. Hogrefe, 2017. doi: 10.1024/85797-000.

[51] Friston, K. J., Holmes, A. P., Worsley, K. J., Poline, J.-P., Frith, C. D. and Frackowiak, R. S. J. ‘Statistical parametric maps in functional imaging: A general linear approach’. In: Human Brain Mapping 2.4 (1994), pp. 189–210. doi: 10.1002/hbm.460020402. eprint: https://onlinelibrary.wiley.com/doi/pdf/10.1002/hbm.460020402. url: https://onlinelibrary.wiley.com/doi/abs/10.1002/hbm.460020402.

[52] Lund, T. E., Nørgaard, M. D., Rostrup, E., Rowe, J. B. and Paulson, O. B. ‘Motion or activity: their role in intra- and inter-subject variation in fMRI’. In: NeuroImage 26.3 (2005), pp. 960–964. issn: 1053-8119. doi: 10.1016/j.neuroimage.2005.02.021. url: https://www.sciencedirect.com/science/article/pii/S1053811905001266.

[53] Siegel, J. S., Power, J. D., Dubis, J. W., Vogel, A. C., Church, J. A., Schlaggar, B. L. et al. ‘Statistical improvements in functional magnetic resonance imaging analyses produced by censoring high-motion data points’. In: Human Brain Mapping 35.5 (2014), pp. 1981–1996. doi: 10.1002/hbm.22307. eprint: https://onlinelibrary.wiley.com/doi/pdf/10.1002/hbm.22307. url: https://onlinelibrary.wiley.com/doi/abs/10.1002/hbm.22307.

[54] Carter, C. S. and Van Veen, V. ‘Anterior cingulate cortex and conflict detection: an update of theory and data’. In: *Cognitive, Affective*, & Behavioral Neuroscience 7.4 (2007), pp. 367–379. doi: 10.3758/CABN.7.4.367.

[55] Kerns, J. G., Cohen, J. D., MacDonald, A. W., Cho, R. Y., Stenger, V. A. and Carter, C. S. ‘Anterior Cingulate Conflict Monitoring and Adjustments in Control’. In: Science 303.5660 (2004), pp. 1023–1026. doi: 10.1126/science.1089910. eprint: https://www.science.org/doi/pdf/10.1126/science.1089910. url: https://www.science.org/doi/abs/10.1126/science.1089910.

[56] Botvinick, M. M., Carter, C. S., Braver, T. S., Barch, D. M. and Cohen, J. D. ‘Conflict monitoring and cognitive control’. In: Psychological Review 108.3 (2001), pp. 624–652. doi: 10.1037/0033-295X.108.3.624.

[57] Peters, S. K., Dunlop, K. and Downar, J. ‘Cortico-Striatal-Thalamic Loop Circuits of the Salience Network: A Central Pathway in Psychiatric Disease and Treatment’. In: Frontiers in Systems Neuroscience 10, 104 (2016). doi: 10.3389/fnsys.2016.00104.

[58] Haber, S. N. ‘Corticostriatal circuitry’. In: Dialogues in Clinical Neuroscience 18.1 (2016), pp. 7–21. doi: 10.31887/DCNS.2016.18.1/shaber.

[59] Weinstein, A. M. ‘Reward, motivation and brain imaging in human healthy participants – A narrative review’. In: Frontiers in Behavioral Neuroscience 17.1123733 (2023), pp. 1–15. issn: 1662-5153. doi: 10.3389/fnbeh.2023.1123733. url: https://www.frontiersin.org/journals/behavioral-neuroscience/articles/10.3389/fnbeh.2023%20.1123733.

[60] Ridderinkhof, K. R., Ullsperger, M., Crone, E. A. and Nieuwenhuis, S. ‘The Role of the Medial Frontal Cortex in Cognitive Control’. In: Science 306.5695 (2004), pp. 443–447. doi: 10.1126/science.1100301. eprint: https://www.science.org/doi/pdf/10.1126/science.1100301. url: https://www.science.org/doi/abs/10.1126/science.1100301.

[61] Domenech, P. and Koechlin, E. ‘Executive control and decision-making in the prefrontal cortex’. In: Current Opinion in Behavioral Sciences 1 (2015). Cognitive control, pp. 101–106. issn: 2352-1546. doi: 10.1016/j.cobeha.2014.10.007. url: https://www.sciencedirect.com/science/article/pii/S2352154614000278.

[62] Smith, J. L., Mattick, R. P., Jamadar, S. D. and Iredale, J. M. ‘Deficits in behavioural inhibition in substance abuse and addiction: A meta-analysis’. In: Drug and Alcohol Dependence 145 (2014), pp. 1–33. doi: 10.1016/j.drugalcdep.2014.08.009. url: https://www.sciencedirect.com/science/article/pii/S0376871614010370.

[63] Wilcox, C. E., Dekonenko, C. J., Mayer, A. R., Bogenschutz, M. P. and Turner, J. A. ‘Cognitive control in alcohol use disorder: deficits and clinical relevance’. In: Reviews in the Neurosciences 25.1 (2013), pp. 1–24. doi: 10.1515/revneuro-2013-0054.

[64] Dinur-Klein, L., Kertzman, S., Rosenberg, O., Kotler, M., Zangen, A. and Dannon, bibinitperiod. ‘Response inhibition and sustained and attention in heavy smokers versus non-smokers’. In: Israel Journal of Psychiatry 51.4 (2014), pp. 240–247.

[65] Luijten, M., Littel, M. and Franken, I. H. A. ‘Deficits in Inhibitory Control in Smokers During a Go/NoGo Task: An Investigation Using Event-Related Brain Potentials’. In: PLOS ONE 6.4 (Apr. 2011), pp. 1–7. doi: 10.1371/journal.pone.0018898. url: 10.1371/journal.pone.0018898.

[66] Nestor, L., McCabe, E., Jones, J., Clancy, L. and Garavan, H. ‘Differences in “bottom-up” and “top-down” neural activity in current and former cigarette smokers: Evidence for neural substrates which may promote nicotine abstinence through increased cognitive control’. In: NeuroImage 56.4 (2011), pp. 2258–2275. issn: 1053-8119. doi: 10.1016/j.neuroimage.2011.03.054. url: https://www.sciencedirect.com/science/article/pii/S1053811911003338.

[67] Dinn, W. M., Aycicegi, A. and Harris, C. L. ‘Cigarette smoking in a student sample: Neurocognitive and clinical correlates’. In: Addictive Behaviors 29.1 (2004), pp. 107–126. doi: 10.1016/j.addbeh.2003.07.001.

[68] Galván, A., Poldrack, R. A., Baker, C. M., McGlennen, K. M. and London, E. D. ‘Neural correlates of response inhibition and cigarette smoking in late adolescence’. In: Neuropsychopharmacology 36.5 (2011), pp. 970–978. doi: doi.org/10.1038/npp.2010.235.

[69] Weywadt, C. R., Kiehl, K. A. and Claus, E. D. ‘Neural correlates of response inhibition in current and former smokers’. In: Behavioural Brain Research 319 (2017), pp. 207–218. issn: 0166-4328. doi: 10.1016/j.bbr.2016.11.030. url: https://www.sciencedirect.com/science/article/pii/S0166432816310877.

[70] Carim-Todd, L., Mitchell, S. H. and Oken, B. S. ‘Impulsivity and Stress Response in Nondependent Smokers (Tobacco Chippers) in Comparison to Heavy Smokers and Nonsmokers’. In: Nicotine & Tobacco Research 18.5 (2016), pp. 547–556. doi: 10.1093/ntr/ntv210.

[71] Domier, C. P., Monterosso, J. R., Brody, A. L., Simon, S. L., Mendrek, A., Olmstead, R. et al. ‘Effects of cigarette smoking and abstinence on Stroop task performance’. In: Psychopharmacology 195 (2007), pp. 1–9. doi: 10.1007/s00213-007-0869-x.

[72] Lesage, E., Sutherland, M. T., Ross, T. J., Salmeron, B. J. and Stein, E. A. ‘Nicotine dependence (trait) and acute nicotinic stimulation (state) modulate attention but not inhibitory control: converging fMRI evidence from Go–Nogo and Flanker tasks’. In: Neuropsychopharmacology 45.5 (2020), pp. 857–865. doi: 10.1038/s41386-020-0623-1.

[73] Charles-Walsh, K., Furlong, L., Munro, D. G. and Hester, R. ‘Inhibitory control dysfunction in nicotine dependence and the influence of short-term abstinence’. In: Drug and Alcohol Dependence 143 (2014), pp. 81–86. doi: 10.1016/j.drugalcdep.2014.07.008.

[74] Wiers, R. W., Pan, T., Dessel, P. van, Rinck, M. and Lindenmeyer, J. ‘Approach-Bias Retraining and Other Training Interventions as Add-On in the Treatment of AUD Patients’. In: Behavioral Neuroscience of Alcohol Addiction: Translational Studies and Human Phenotypes. Ed. by Sommer, W. H. and Spanagel, R. Cham: Springer Nature Switzerland, 2025, pp. 641–687. isbn: 978-3-032-01585-3. doi: 10.1007/7854_2023_421.

[75] Boffo, M., Zerhouni, O., Gronau, Q. F., van Beek, R. J. J., Nikolaou, K., Marsman, M. et al. ‘Cognitive Bias Modification for Behavior Change in Alcohol and Smoking Addiction: Bayesian Meta-Analysis of Individual Participant Data’. In: Neuropsychology Review 29.1 (2019), pp. 52–78. issn: 1573-6660. doi: 10.1007/s11065-018-9386-4.

[76] Cristea, I. A., Kok, R. N. and Cuijpers, P. ‘The Effectiveness of Cognitive Bias Modification Interventions for Substance Addictions: A Meta-Analysis’. In: PLOS ONE 11.9 (Sept. 2016), pp. 1–19. doi: 10.1371/journal.pone.0162226. url: 10.1371/journal.pone.0162226.

[77] Zhang, M. and Peng, Y. ‘Anterior insula and dorsal anterior cingulate cortex as a hub of self-regulation: combining activation likelihood estimation meta-analysis and meta-analytic connectivity modeling analysis’. In: Brain Structure and Function 228.5 (2023), pp. 1329–1345. issn: 1863-2661. doi: 10.1007/s00429-023-02652-9. url: 10.1007/s00429-023-02652-9.

[78] Liu, X., Hairston, J., Schrier, M. and Fan, J. ‘Common and distinct networks underlying reward valence and processing stages: A meta-analysis of functional neuroimaging studies’. In: Neuroscience & Biobehavioral Reviews 35.5 (2011), pp. 1219–1236. issn: 0149-7634. doi: 10.1016/j.neubiorev.2010.12.012. url: https://www.sciencedirect.com/science/article/pii/S0149763410002137.

[79] Tzourio-Mazoyer, N., Landeau, B., Papathanassiou, D., Crivello, F., Etard, O., Delcroix, N. et al. ‘Automated Anatomical Labeling of Activations in SPM Using a Macroscopic Anatomical Parcellation of the MNI MRI Single-Subject Brain’. In: NeuroImage 15.1 (2002), pp. 273–289. issn: 1053-8119. doi: 10.1006/nimg.2001.0978. url: https://www.sciencedirect.com/science/article/pii/S1053811901909784.

[80] Moeller, S., Yacoub, E., Olman, C. A., Auerbach, E., Strupp, J., Harel, N. et al. ‘Multiband multislice GE-EPI at 7 tesla, with 16-fold acceleration using partial parallel imaging with application to high spatial and temporal whole-brain fMRI’. In: Magnetic Resonance in Medicine 63.5 (2010), pp. 1144–1153. doi: 10.1002/mrm.22361. eprint: https://onlinelibrary.wiley.com/doi/pdf/10.1002/mrm.22361. url: https://onlinelibrary.wiley.com/doi/abs/10.1002/mrm.22361.

[81] Bates, D., Mächler, M., Bolker, B. and Walker, S. ‘Fitting Linear Mixed-Effects Models Using lme4’. In: Journal of Statistical Software 67.1 (2015), pp. 1–48. doi: 10.18637/jss.v067.i01. url: https://www.jstatsoft.org/index.php/jss/article/view/v067i01.

