## Supplementary Material for "Interference Between Pavlovian Cues and Instrumental Control: A Stratified Comparison Across Alcohol Use Disorder and Smoking"

### Supplementary Material S1

#### QFQ Smoking

The quantity and frequency of tobacco use were assessed at each time point using a novel quantity-frequency questionnaire. The questionnaire was administered as a structured interview by trained experimenters and covered the use of cigarettes, e-cigarettes, and other tobacco products over the last three months. After an initial question targeting the types of tobacco products consumed during this period, the term tobacco was used as a collective term referring to all tobacco products used by the participant.

**Table S1.1**

*English Translation of the Quantity-Frequency Questionnaire for Smoking*

| n° | Type | Item |
|---|---|---|
| 1 | MC | What did you smoke? ('*Cigarettes*', '*Cigarillos*', '*Cigars*', '*Hookah (Shisha)*', '*E-cigarette*', '*Tobacco heating-systems*', '*Snuff*', '*Other tobacco product*') |
| 2a | ON | On how many days in the last 3 months did you smoke at least 1 cigarette/use an e-cigarette/tobacco product at least once? |
| 2b | SC | What format were the days given in the previous question? ('*Reported over the past 3 months*', '*Reported per week over the past 3 months*') |
| 3 | ON | When you were smoking the most over the last 3 months, how many cigarettes did you smoke on average each day/how often did you use an e-cigarette on average? |
| 4 | SC | Have you ever taken part in a smoking cessation programme? For example, using nicotine patches, behavioural therapy or something similar? ('*Yes*', '*No*') |
| 5a | SC | Have you ever smoked regularly in your life, that is, have you smoked at least one cigarette (or another tobacco product) a day over a period of 4 weeks or longer? ('*Yes*', '*No*') |
| (5b) [a] | ON | When was the first time you smoked at least one cigarette a day or used an e-cigarette every day for a period of more than 4 weeks or longer? |
| (5c) [a] | ON | When was the first time you smoked at least one cigarette a day or used an e-cigarette every day for a period of more than 4 weeks or longer? |

*Note.* MC = multiple choice; ON = open numeric; SC = single choice.

[a] Displayed only if question 5a was answered 'No'.

### Supplementary Material S2

### Assessment of Task Understanding & Participant Exclusions

Task understanding was assessed based on participants' behavior during the PIT phase (Part 3), the query trials (Part 4), and the debriefing questionnaire.

#### Query Trials

During the query trials (Part 4), participants were forced to choose between pairs of Pavlovian CSs. In each trial, two compound cues from Part 2 were sequentially presented on opposite sides of the screen. Participants were instructed to choose between the two options based on their subjective preference using the left and right arrow keys. Each CS pairing was presented 6 times, resulting in a total of 60 query trials in randomized order.

To assess whether participants had learned the relative valence of money-related Pavlovian CSs during Part 2, we computed how often participants chose a higher-valued Pavlovian CS option over a lower-valued CS option (e.g., '+10€' CS over '-10€' CS; preference score). As illustrated in Figure S2.1, participants with a preference score < 50% across all money-related trials, indicating insufficient awareness of the Pavlovian CS's relative valence, were excluded from the analysis.

#### PIT phase & Debriefing Questionnaire

Participants were also excluded when they exhibited fully Pavlovian CS-dependent behavior (100% ER in incongruent and 0% ER in congruent trials) while agreeing to a task-related statement that fractal-like images (i.e., Pavlovian CSs) predicted the valence of the instrumental shells in the debriefing questionnaire (i.e., 'The abstract backgrounds indicated whether a shell was good or bad.'). In these cases, the observed maximum PIT interference may have reflected an erroneous belief in a CS-shell valence contingency rather than genuine PIT.

Finally, participants who performed below chance level during congruent PIT trials (ER > 50%), in which the Pavlovian CS and the instrumental stimulus had the same valence, thereby precluding PIT-related conflicts, were also excluded.

**Exclusions**

Based on these criteria, 39 participants were excluded: 4 because they exhibited fully Pavlovian CS-dependent behavior while simultaneously reporting that fractal-like images (i.e., Pavlovian CSs) predicted the valence of the instrumental shells in the debriefing questionnaire, 14 because they performed below chance level in the congruent condition, and 21 because they selected the lower-value option in more than 50% of the query trials.

**Figure S2.1**

*Mean Error Rates Across Congruity Conditions for Participants With High (>50%) and Low (<50%) Query-Trial Preference Scores*

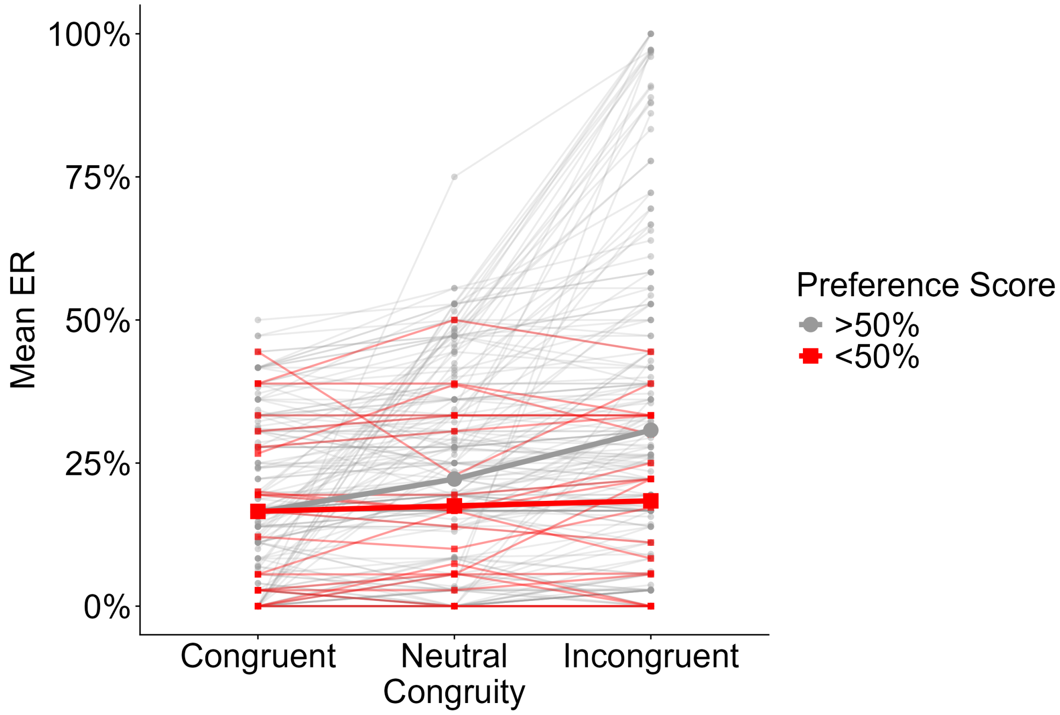


*Note.* Each small dot represents a participant's individual mean error rate (ER) in congruent, neutral, or incongruent PIT trials, with thin lines illustrating within-person ER changes across all three congruity conditions. Thicker dots and lines represent group-level means and congruity-related changes for participants who chose the higher-valued option in more than 50% of the query trials (preference score >50%; gray) and participants who did not choose the higher-valued option in more than 50% of the query trials (preference score <50%; red). Participants with a preference score >50% showed the expected PIT interference effect, with the mean ER increasing from the congruent to the incongruent condition. Participants with a preference score <50% showed comparable ERs across all three congruity conditions, at a level similar to the mean of the congruent condition for the >50% preference score group. Because a preference score <50% indicates an insufficient acquisition of the Pavlovian cue–outcome associations during the Pavlovian conditioning phase, these participants were excluded from all subsequent analyses.

### Supplementary Material S3

### MRI Acquisition & Preprocessing

#### MRI Acquisition

All MRI data were recorded on a 3T Siemens MAGNETOM® Tim Trio (Siemens Healthineers AG, Forchheim, Germany) using a 32-channel head coil. First, high-resolution structural images were acquired using a T1-weighted magnetization-prepared rapid gradient-echo sequence with repetition time (TR) = 2000 ms, echo time (TE) = 2.01 ms, inversion time (TI) = 880 ms, bandwidth = 240 Hz/pixel, and flip-angle (FA) = 8°. A total of 208 sagittal slices were obtained with slice thickness = 1 mm and field of view (FOV) = 256 mm, yielding a voxel size of 1 x 1 x 1 $mm^3$. To correct for geometric distortions during functional imaging as a result of static field inhomogeneities, a field map was then acquired using a Siemens dual gradient-echo sequence with 64 interleavingly acquired slices, voxel size = 2.4 x 2.4 x 2.4 $mm^3$, TR = 698 ms, TE1 = 5.19, TE2 = 7.65 ms, bandwidth = 279 Hz/pixel, FOV = 210 mm. Functional images were acquired during the PIT phase using a multi-band T2*-weighted gradient-echoplanar imaging (EPI) sequence [80] with slice acceleration factor = 6, TR = 869 ms, TE = 38 ms, and FA = 58°. For each scan, 60 slices covering the whole brain were acquired interleavingly with slice thickness = 2.4 mm, bandwidth = 1832 Hz/pixel, and FOV = 210 mm, yielding a voxel size of 2.4 x 2.4 x 2.4 $mm^3$.

#### MRI Preprocessing

The imaging data were preprocessed using SPM12 (Wellcome Department of Cognitive Neurology, Institute of Neurology, London, United Kingdom). Functional images were slice-time corrected using the middle slice as reference, realigned to the first image and then the mean image, and corrected for geometric distortions using voxel-displacement maps derived from individual field maps. Structural images were co-registered to the mean EPI images and segmented into gray and white matter. Using the same T1-derived transformation obtained during segmentation, functional and structural images were normalized to the standard Montreal Neurological Institute space (ICBM 152 MNI template) and subsequently resampled to their respective original voxel size. In a final step, EPI images were smoothed using an 8-mm full-width-at-half-maximum Gaussian kernel.

### Supplementary Material S4

### Preprocessing of Joystick Data & Task Manipulation Check

Prior to the main behavioral analysis, the raw joystick data from the PIT phase were preprocessed using MATLAB R2024B (version 24.2.0.2806996; The MathWorks, Inc., 2022). Preprocessing included the trial-by-trial computation of the initial joystick position, the peak velocity of the joystick movement, and the correctness of the action taken. During this step, a subset of trials (4.40%) was identified in which the joystick failed to register as being in the neutral position at trial onset. To minimize the potential confounding by this technical artifact, trials with an initial joystick position exceeding 20% of the full movement range were excluded. Participants with more than 40% of affected trials were excluded entirely ($n = 4$).

We then conducted an initial task manipulation check to assess whether participants committed more errors during incongruent PIT trials, characterized by a conflict between a Pavlovian CS and the correct instrumental response, than during congruent trials without such conflict. PIT trials were therefore classified according to congruity before computing the mean error rate (ER) for each congruity condition. The ERs of both congruity conditions were then compared using a paired $t$-test. Based on previous evidence of substantial PIT interference across all participants [13], we expected higher ERs across incongruent PIT trials than congruent PIT trials. Similar to Chen et al. [13], we observed a substantial PIT interference effect across the entire sample ($M$ = 14.28%, $SD$ = 28.54%), $V = 10384$, $p$(one-sided) $< .001$, $r = .56$.

**Figure S4.1**

*PIT Interference Across the Entire Sample*

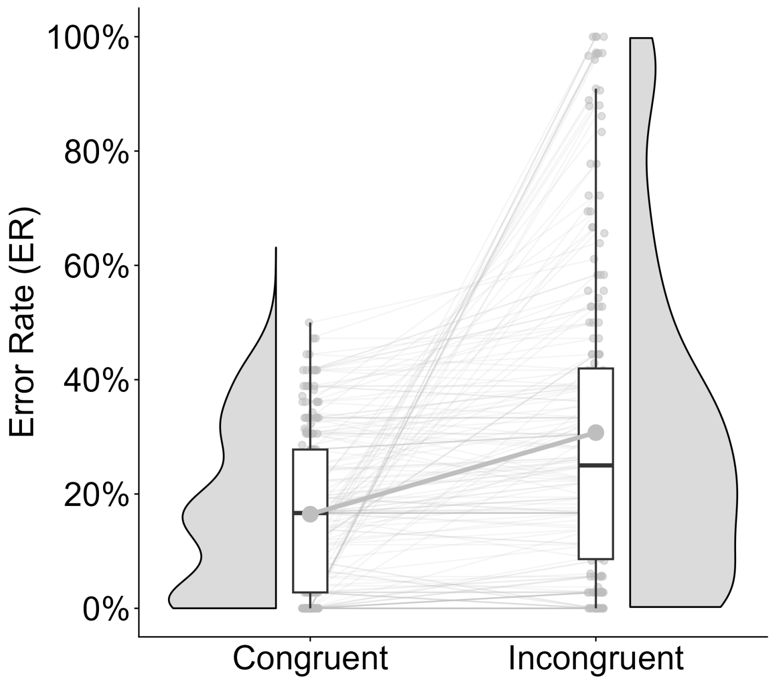


*Note.* Each dot represents an individual participant, with thin lines illustrating the individual changes of the mean error rate from the congruent to the incongruent condition and the thick line illustrating the mean change across all participants. The half-violin plots and boxplots illustrate the distribution of the mean ERs within the congruent and incongruent conditions.

### Supplementary Material S5

### Motivational PIT Analysis

Given the previous reports of a more pronounced Pavlovian CS-dependent invigoration and suppression of instrumental approach behavior in risky drinkers [9, 10] and detoxified alcohol-dependent inpatients [11], we also conducted an exploratory analysis of motivational PIT in the present sample.

#### Statistical Analysis

Motivational PIT was operationalized as the trial-by-trial modulation of the mean joystick velocity by money-related Pavlovian CSs. The mean velocity was computed for each trial by dividing the distance from movement onset to decision threshold by movement time and analyzed using linear mixed-effect models (lmer), implemented in the *lme4* package [81]. In each model, the mean velocity was predicted by Pavlovian CS valence (-1 = '-10€', 0 = 'neutral', 1 = '+10€'), shell valence (0.5 = 'good', -0.5 = 'bad'), AUD group (-0.5 = 'non-AUD', 0.5 = 'AUD'), smoking status group (mean-centered), and all higher-order and lower-order interactions. To determine the optimal random-effects structure, models of increasing complexity were fitted using maximum likelihood estimation and compared using likelihood ratio tests, Akaike's information criterion (AIC), and Bayesian information criterion (BIC). As summarized in Table S5.1, the model including random effects for Pavlovian CS valence and shell valence, along with random intercepts for individual shell stimuli, provided the best overall fit and was therefore selected as the final model for the motivational PIT analysis. Consistent with the behavioral PIT interference analysis, an extended version of this model was fitted, including age and gender as additive covariates.

To assess the association between motivational PIT and PIT interference, we extracted participant-specific random slopes for Pavlovian CSs from the final lmer model and correlated these estimates with $\Delta ER$ using Pearson's correlation coefficient [cf. 13].

We also tested the dimensional associations between motivational PIT and continuous substance-related measures. For this, the winning model was refitted after replacing the fixed effects for AUD, smoking status, and AUD x smoking status

**Table S5.1**

*Model Comparison*

| | npar | AIC | BIC | logLik | $\chi^2$ | df | p |
|---|---|---|---|---|---|---|---|
| **(A) Determine if random slopes for PavCS and InsCond improve model fit** | | | | | | | |
| Model 1: | *+ (1 / subj)* | | | | | | |
| Model 2: | *+ (1 + PavCS + InsCond / subj)* | | | | | | |
| Model 1: | 18 | 239214 | 239356 | -119589 | | | |
| Model 2: | 23 | 234066 | 234249 | -117010 | 5157.378 | 5 | **<.001** |
| **(B1) Determine if random effects for shell stimuli improve model fit** | | | | | | | |
| Model 2: | *+ (1 + PavCS + InsCond / subj)* | | | | | | |
| Model 3: | *+ (1 + PavCS + InsCond / subj) + (1 / shell)* | | | | | | |
| Model 2: | 23 | 234066 | 234249 | -117010 | | | |
| Model 3: | 24 | 233973 | 234164 | -116963 | 94.817 | 1 | **<.001** |
| **(B2) Determine if random effects for audiovisual compound cues improve model fit** | | | | | | | |
| Model 2: | *+ (1 + PavCS + InsCond / subj)* | | | | | | |
| Model 4: | *+ (1 + PavCS + InsCond / subj) + (1 / compound cue)* | | | | | | |
| Model 2: | 23 | 234066 | 234249 | -117010 | | | |
| Model 4: | 24 | 234066 | 234257 | -117009 | 1.7614 | 1 | .184 |
| **(C) Determine if random effects for shell stimuli and compound cues improve model fit** | | | | | | | |
| Model 3: | *+ (1 + PavCS + InsCond / subj) + (1 / shell)* | | | | | | |
| Model 5: | *+ (1 + PavCS + InsCond / subj) + (1 / shell) + (1 / compound cue)* | | | | | | |
| Model 3: | 24 | 233973 | 234164 | -116963 | | | |
| Model 5: | 25 | 233974 | 234172 | -116962 | 1.7738 | 1 | .183 |

*Note.* npar = number of parameters; AIC = Akaike information criterion; BIC = Bayesian information criterion; logLik = log-likelihood. The base model (Model 1) predicted the trial-by-trial mean velocity (MV) from the fixed effects of Pavlovian conditioned stimuli (PavCS), instrumental condition (InsCond), alcohol use disorder (AUD), smoking status, and all lower- and higher-order interactions among these factors, with a random intercept for participants (subj): MV ~ PavCS * InsCond * AUD * SmokingStatus + (1|subj)

interaction with AUDIT or FTND total score (disorder severity), AUDIT-C (overall drinking), the number of drinking or smoking days per week (consumption frequency), the average amount of daily alcohol or tobacco intake (consumption quantity), CAS-A total or QSU-G mean score (craving).

### Results

As expected, the lmer revealed a significant modulation of the PV by Pavlovian CSs, $\beta = 11.96$, $SE = 1.73$, $p$(two-sided) $< .001$, 95% CI [6.57, 15.35], indicating substantial motivational PIT across all participants (see Figure S5.1). Unexpectedly, however, the magnitude of the motivational PIT effect did not differ between the two AUD groups, $\beta = 4.40$, $SE = 3.46$, $p$(two-sided) = .203, 95% CI [-2.38, 11.18], nor did it change as a function of increasing smoking status, $\beta = -3.01$, $SE = 2.17$, $p$(two-sided) = .166, 95% CI [-7.26, 1.25]. There was also no evidence for a significant interaction between AUD and smoking status with respect to motivational PIT, $\beta$ = -2.58, $SE$ = 3.34, $p$(two-sided) = .551, 95% CI [-11.09, 5.92]. Including age and gender as covariates did not alter the overall pattern of the results and revealed no significant age or gender effect on motivational PIT (see Table S5.2).

**Figure S5.1**

*Motivational PIT Across Sample*

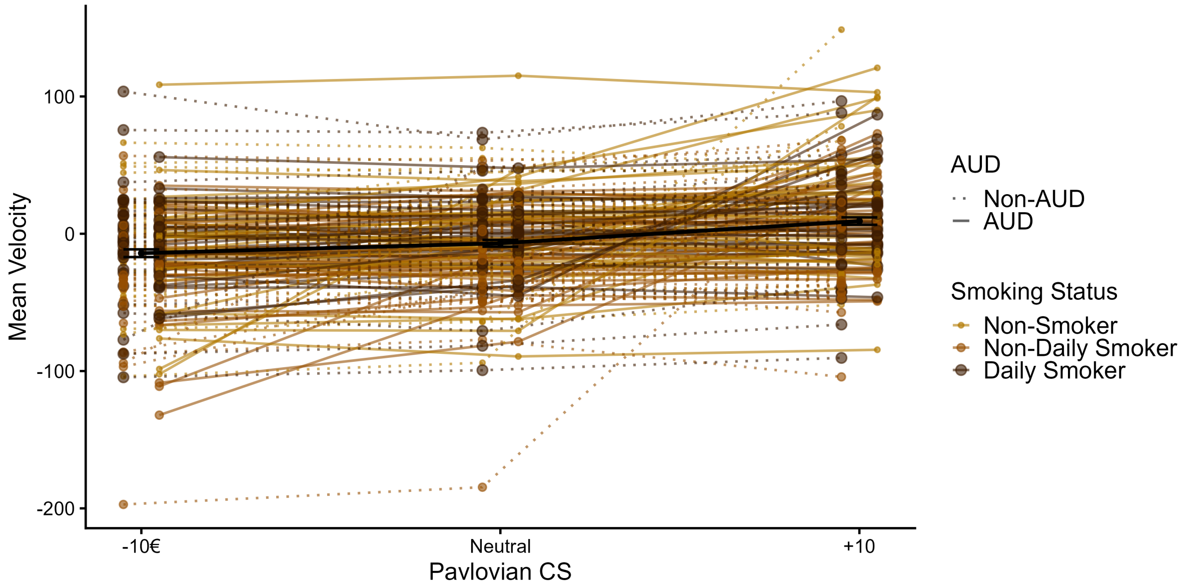


*Note.* Smaller colored dots and triangles represent participants' individual mean joystick velocity (MV) during the PIT phase, with lines illustrating the within-person MV change across the three Pavlovian CS conditions (-10€, neutral, +10€). The thicker dots and lines represent the modulation of MV at group level, with error bars illustrating the between-subject standard deviation within each Pavlovian CS condition.

The correlation between motivational PIT and PIT interference was highly significant and positive, $r = .92$, $p$(two-sided) $< .001$, indicating that individuals with a Pavlovian CS-dependent modulation of the response vigor were also more likely to

**Table S5.2**

*Results From Age- and Gender-Adjusted lmer on Motivational PIT*

| Effect | Estimate | *SE* | 95% CI | | *p* |
|---|---|---|---|---|---|
| | | | LL | UL | |
| **Main Effects** | | | | | |
| (Intercept) | -8.69 | 7.58 | -23.54 | 6.16 | .251 |
| Pav CS | 11.96 | 1.73 | 8.57 | 15.35 | **<.001** |
| Ins Cond | 106.00 | 5.44 | 95.34 | 116.66 | **<.001** |
| AUD | 0.80 | 4.31 | -7.65 | 9.25 | .853 |
| Smoking Status | -1.64 | 2.67 | -6.87 | 3.59 | .538 |
| Age | 0.17 | 0.22 | -0.26 | 0.59 | .444 |
| Gender | -1.18 | 4.27 | -9.56 | 7.19 | .782 |
| **Two-Way Interaction Effects** | | | | | |
| Pav CS x Ins Cond | 0.75 | 1.18 | -1.56 | 3.07 | .523 |
| Pav CS x AUD | 4.40 | 3.46 | -2.38 | 11.18 | .203 |
| Pav CS x Smoking Status | -3.01 | 2.17 | -7.26 | 1.25 | .166 |
| Ins Cond x AUD | 4.31 | 10.88 | -17.01 | 25.62 | .692 |
| Ins Cond x Smoking Status | -1.62 | 6.82 | -14.98 | 11.75 | .812 |
| AUD x Smoking Status | 2.28 | 5.40 | -8.31 | 12.87 | .673 |
| **Three-Way Interaction Effects** | | | | | |
| Pav CS x Ins Cond x AUD | -1.64 | 2.36 | -6.27 | 2.98 | .486 |
| Pav CS x Ins Cond x Smoking Status | -0.17 | 1.48 | -3.08 | 2.73 | .906 |
| Pav CS x AUD x Smoking Status | -2.58 | 4.34 | -11.09 | 5.92 | .551 |
| Ins Cond x AUD x Smoking Status | -2.61 | 13.64 | -29.34 | 24.12 | .848 |
| **Four-Way Interaction Effects** | | | | | |
| Pav CS x Ins Cond x AUD x Smoking | -3.43 | 2.96 | -9.23 | 2.37 | .247 |

*Note.* $N = 200$, total number of observations = 20543. SE = standard error; CI = confidence interval; LL = lower limit; UL = upper limit; Pav CS = Pavlovian conditioned stimulus; Ins Cond = Instrumental condition (shell valence); AUD = Alcohol Use Disorder.

commit errors when the Pavlovian CSs conflicted with the correct instrumental response.

With respect to the continuous substance-related measures, only drinking frequency significantly moderated the effect of Pavlovian CSs on mean velocity, indicating that more frequent alcohol consumption was associated with stronger motivational PIT effects. As summarized in Table S5.3, none of the other variables examined showed a significant moderating effect. It should be noted, however, that both TUD severity (FTND total score), $\beta = -6.89$, $SE = 2.87$, $p$(two-sided) = .016, 95% CI [-12.50, -1.27], and nicotine craving (QSU-G score), $\beta = -13.28$, $SE = 5.19$, *p*(two-sided) = .010, 95% CI [-23.44, -3.11], significantly moderated the effects of instrumental condition on mean velocity. These findings indicate that greater nicotine dependence and craving were associated with a reduced influence of shell valence on mean velocity.

**Table S5.3**

*Dimensional Associations Between Motivational PIT and Continuous Substance-Related Measures*

| | Estimate | *SE* | 95% CI | | *p* |
|---|---|---|---|---|---|
| | | | LL | UL | |
| **Disorder Severity** | | | | | |
| AUDIT Total Score [a] | 0.002 | 0.25 | -0.49 | 0.50 | .993 |
| FTND Total Score [b] | -0.79 | 0.89 | -2.54 | 0.96 | .375 |
| **Consumption** | | | | | |
| Alcohol | | | | | |
| AUDIT-C [a] | 0.763 | 0.66 | -0.53 | 2.06 | .247 |
| Quantity (g/day) [c] | 0.03 | 0.04 | -0.06 | 0.11 | .543 |
| Frequency (days/week) [c] | 1.71 | 0.83 | 0.09 | 3.33 | **.039** |
| Tobacco | | | | | |
| Quantity (units/day) [d] | -0.28 | 0.26 | -0.78 | 0.23 | .287 |
| Frequency (days/week) [e] | -0.64 | 0.56 | -1.74 | 0.45 | .250 |
| **Craving** | | | | | |
| CAS-A [a] | 0.12 | 0.17 | -0.21 | 0.45 | .491 |
| QSU-G [f] | -1.08 | 1.82 | -4.63 | 2.48 | .553 |

*Note.* PIT = Pavlovian-to-instrumental transfer; SE = standard error; CI = confidence interval; LL = lower limit; UL = upper limit; AUDIT = Alcohol Use Disorders Identification Test; FTND = Fagerström Test for Nicotine Dependence; AUDIT-C = Alcohol Use Disorders Identification Test - Consumption; g = grams; CAS-A = Craving Automated Scale for Alcohol; QSU-G = Questionnaire on Smoking Urges - German.
[a] $N$ = 185, total number of observations = 18999. [b] $N$ = 167, total number of observations = 17140. [c] $N$ = 199, total number of observations = 20435. [d] $N$ = 199, total number of observations = 20437. [e] $N$ = 200, total number of observations = 20543. [f] $N$ = 93, total number of observations = 9474.

### Supplementary Material S6

#### Additional Information on the Stratified Sample

Figure S6.1 illustrates the distribution of the AUD and TUD criteria across the entire sample. According to the stratification design, none of the participants in the non-AUD group and all participants in the AUD group surpassed the diagnostic threshold of two DSM-5 criteria for AUD. With 76%, the majority of participants in the AUD group met fewer than six AUD criteria and can therefore be classified as having mild to moderate AUD. With respect to smoking, 42.1% of the occasional smokers and 86.2% of the daily smokers meet at least two of the DSM-5 criteria for a TUD and could thereby be classified as clinically relevant cases. In the non-smoker group who reported no smoking in the last three months before the study, one participant met two DSM-5 criteria over a 12-month period during SCID, thereby surpassing the diagnostic threshold for TUD.

Table S6.1 provides a more detailed description of the sample by summarizing sociodemographic information and continuous alcohol- and smoking-related measures included in this study for all six stratification groups.

**Figure S6.1**

*Distribution of AUD and TUD Criteria Across Sample*

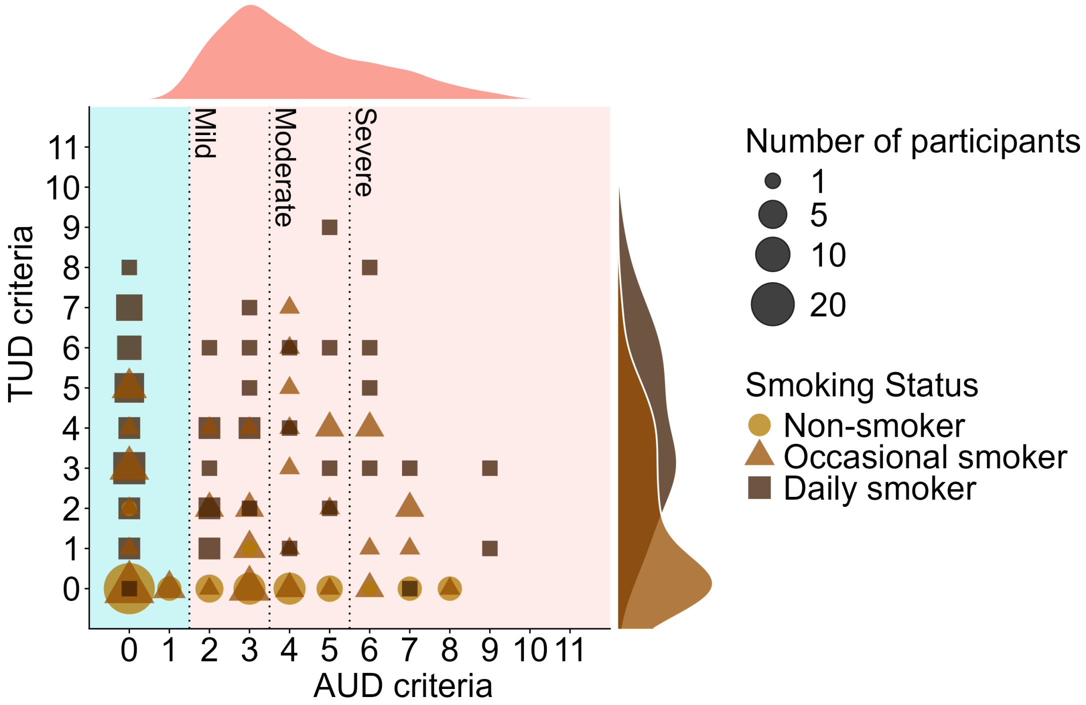


*Note.* Each bubble represents a group of participants fullfilling the same number of AUD and TUD criteria, with the size of the bubble indicating the group and the shape indicating smoking status. The distribution plots next to the bubble plot represent the distributions of AUD criteria (top) and TUD criteria (right) for participants classified as 'AUD' (top) or 'occasional smoker' or 'daily smoker' (right).

**Table S6.1**

*Sample Descriptives by AUD and Smoking Status*

| | Non-AUD | | | AUD | | |
|---|---|---|---|---|---|---|
| | Non-Smoker | Occasional Smoker | Daily Smoker | Non-Smoker | Occasional Smoker | Daily Smoker |
| **Sociodemographic Characteristics** | | | | | | |
| N | 37 | 34 | 29 | 33 | 38 | 29 |
| Age | 28.3 (±8.8) | 30.0 (±8.2) | 31.3 (±8.2) | 37.6 (±12.6) | 29.6 (±10.0) | 32.8 (±9.0) |
| Gender (♀) | 43.2% | 41.2% | 41.4% | 42.4% | 39.5% | 37.9% |
| Education | | | | | | |
| Low | 2.7% | 0.0% | 6.9% | 15.2% | 0.0% | 6.9% |
| Middle | 5.4% | 11.8% | 13.8% | 24.2% | 5.3% | 20.7% |
| High | 83.8% | 82.4% | 65.5% | 57.6% | 86.8% | 62.1% |
| **Alcohol-related measures** | | | | | | |
| AUD criteria | 0.1 (±0.3) | 0.1 (±0.3) | 0.0 (±0.0) | 4.2 (±1.9) | 4.2 (±1.6) | 4.2 (±2.1) |
| AUDIT | 3.0 (±3.1) | 4.5 (±2.8) | 4.7 (±2.9) | 15.0 (±6.0) | 15.3 (±4.0) | 14.3 (±4.8) |
| AUDIT-C | 2.3 (±1.6) | 3.0 (±1.3) | 3.3 (±1.5) | 6.6 (±2.1) | 6.9 (±1.6) | 7.0 (±1.5) |
| Consumption | | | | | | |
| grams/day | 28.1 (±20.9) | 33.7 (±22.5) | 37.4 (±27.5) | 71.7 (±47.9) | 74.6 (±39.0) | 82.3 (±32.9) |
| days/week | 0.8 (±0.9) | 1.2 (±1.3) | 1.4 (±1.4) | 4.3 (±1.8) | 3.9 (±1.5) | 4.1 (±1.7) |
| CAS-A | 1.9 (±4.2) | 2.8 (±4.0) | 4.7 (±7.3) | 17.2 (±9.6) | 15.8 (±9.9) | 15.6 (±9.9) |
| **Smoking-related measures** | | | | | | |
| TUD criteria | 0.1 (±0.3) | 1.6 (±1.9) | 4.2 (±2.1) | 0.0 (±0.2) | 1.8 (±1.9) | 3.8 (±2.3) |
| FTND | 0.0 (±0.0) | 0.3 (±0.7) | 3.5 (±2.5) | 0.1 (±0.7) | 0.2 (±0.5) | 3.0 (±2.6) |
| Consumption | | | | | | |
| units/day | 0.0 (±0.0) | 2.7 (±1.9) | 13.1 (±5.4) | 0.0 (±0.0) | 3.3 (±3.2) | 13.4 (±7.8) |
| days/week | 0.0 (±0.0) | 1.5 (±1.9) | 7.0 (±0.0) | 0.0 (±0.0) | 1.6 (±1.9) | 7.0 (±0.0) |
| QSU-G | / | 1.9 (±1.0) | 3.4 (±1.2) | 2.8 (±0.0) | 2.0 (±1.0) | 3.5 (±1.2) |

*Note.* AUD = Alcohol Use Disorder; AUDIT = AUD Identification Test; AUDIT-C = AUDIT-Consumption; CAS-A = Craving Automated Scale for Alcohol; TUD = Tobacco Use Disorder; QSU-G = Questionnaire on Smoking Urges - German version.

### Supplementary Material S7

### Interaction of AUD and Smoking Status With Respect to Behavioral PIT Interference

**Figure S7.1**

*Modulation of AUD Group Difference in PIT Interference by Smoking Status*

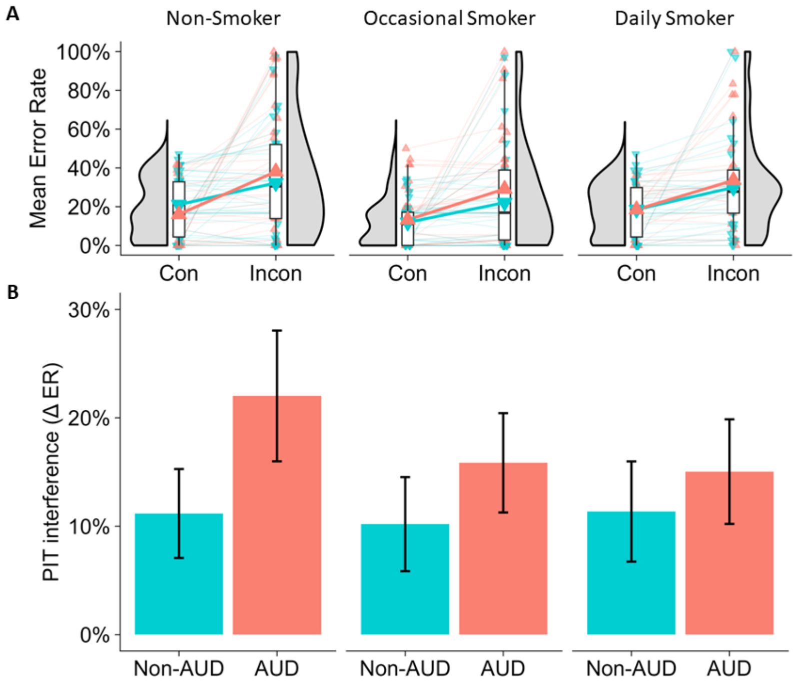


*Note.* Each dot represents an individual participant, with thin lines illustrating the individual changes of the mean error rate from the congruent to the incongruent condition and the thick line illustrating the mean change across all participants. The half-violin plots and boxplots illustrate the distributions of mean ERs across the congruent and incongruent conditions.

### Supplementary Material S8

### Neural PIT Interference

**Table S8.1**

*Association Between Behavioral and Neural PIT Interference in Selected ROIs*

| | $H_{Dir.}$ | $\tau$ | $p$ |
|---|---|---|---|
| **Neural PIT interference (incongruent vs. congruent)** | | | |
| right lPFC | - | -.01 | .401 |
| bilateral dmPFC | - | .01 | .532 |
| bilateral VS | + | -.02 | .622 |
| bilateral amygdala | + | -.08 | .943 |
| ACC | - | .06 | .885 |

*Note.* Kendall's $\tau$ correlations were tested one-sided according to a priori directional hypotheses. $H_{Dir.}$ = hypothesized direction; ROI = Region of Interest; lPFC = lateral prefrontal cortex; dmPFC = dorsomedial prefrontal cortex; VS = ventral striatum.

**Table S8.2**

*Results of Exploratory Whole-Brain Analysis of Neural PIT Interference*

| | | Cluster | | Peak | | | |
|---|---|---|---|---|---|---|---|
| | | $k$ | $p_{FWE}$ | MNI coordinates | | | $t$ |
| | | | | x | y | z | |
| L | Superior frontal gyrus | 1358 | **<.001** | -1 | 22 | 48 | 5.56 |
| R | Angular gyrus | 633 | **<.001** | 54 | -42 | 48 | 4.39 |
| R | Insular cortex | 443 | **.001** | 30 | 20 | -8 | 5.13 |
| L | Insular cortex | 232 | **.018** | -32 | 18 | -10 | 4.84 |
| L | Supramarginal gyrus, posterior division | 116 | .157 | -56 | -57 | 48 | 3.88 |
| R | Cingulate gyrus, posterior division | 99 | .222 | 6 | -26 | 36 | 3.73 |
| R | Middle frontal gyrus | 92 | .256 | 42 | 32 | 33 | 4.18 |
| R | Middle frontal gyrus | 77 | .347 | 42 | 18 | 45 | 3.53 |
| L | Superior frontal gyrus | 65 | .441 | -11 | 13 | 64 | 3.75 |
| R | Frontal pole | 57 | .514 | 25 | 61 | 2 | 3.84 |

*Note.* $k$ = Cluster Size; FWE = Family-wise Error Corrected; MNI = Montreal Neurological Institute. The second-level contrast image for neural PIT interference was subjected to a one-sample $t$-test, with a significance threshold of $p$(uncorrected) $< .001$ and a minimum cluster extent of $k > 50$ voxels.

**Table S8.3**

*AUD x Smoking Status Effects on Neural PIT Interference*

| | $H_{Dir.}$ | Estimate | *SE* | 95% CI | | $p_{FDR}$ |
|---|---|---|---|---|---|---|
| | | | | LL | UL | |
| **Right lPFC** | | | | | | |
| AUD > Non-AUD | - | -.18 | .16 | -.50 | .13 | .345 |
| Smoking Status | - | .13 | .14 | -.13 | .40 | .822 |
| AUD x Smoking Status | +/- | -.10 | .20 | -.49 | .28 | .981 |
| **Bilateral dmPFC** | | | | | | |
| AUD > Non-AUD | - | -.16 | .17 | -.48 | .19 | .348 |
| Smoking Status | - | .06 | .15 | -.24 | .35 | .822 |
| AUD x Smoking Status | +/- | -.09 | .21 | -.52 | .33 | .981 |
| **Bilateral VS** | | | | | | |
| AUD > Non-AUD | + | -.11 | .16 | -.42 | .19 | .816 |
| Smoking Status | + | .17 | .14 | -.11 | .45 | .465 |
| AUD x Smoking Status | +/- | .04 | .20 | -.35 | .45 | .981 |
| **Bilateral amygdala** | | | | | | |
| AUD > Non-AUD | + | -.17 | .19 | -.52 | .21 | .816 |
| Smoking Status | + | .01 | .16 | -.30 | .33 | .822 |
| AUD x Smoking Status | +/- | -.01 | .23 | -.48 | .44 | .981 |

*Note.* $N = 180$. $H_{Dir.}$ = hypothesized direction; SE = standard error; CI = confidence interval (bias-corrected and accelerated); LL = lower limit; UL = upper limit; lPFC = lateral prefrontal cortex; dmPFC = dorsomedial prefrontal cortex; VS = ventral striatum; ACC = anterior cingulate cortex; AUD = Alcohol Use Disorder.
